# Internalization of the host alkaline pH signal in a fungal pathogen

**DOI:** 10.1101/2020.10.19.345280

**Authors:** Hannah E. Brown, Kaila M. Pianalto, Caroline M. Fernandes, Katherine D. Mueller, Maurizio Del Poeta, J. Andrew Alspaugh

## Abstract

The ability for cells to internalize extracellular cues allows them to adapt to novel and stressful environments. This adaptability is especially important for microbial pathogens that must sense and respond to drastic changes when encountering the human host. *Cryptococcus neoformans* is an environmental fungus and opportunistic pathogen that naturally lives in slightly acidic reservoirs, but must adapt to the relative increase in alkalinity in the human host in order to effectively cause disease. The fungal-specific Rim alkaline response signaling pathway effectively converts this extracellular signal into an adaptive cellular response allowing the pathogen to survive in its new environment. The newly identified Rra1 protein, the most upstream component of the *C. neoformans* Rim pathway, is an essential component of this alkaline response. Previous work connected Rra1-mediated signaling to the dynamics of the plasma membrane. Here we identify the specific mechanisms of Rim pathway signaling through detailed studies of the activation of the Rra1 protein. Specifically, we observe that the Rra1 protein is internalized and recycled in a pH-dependent manner, and that this dynamic pattern of localization further depends on specific residues in its C-terminal tail, clathrin-mediated endocytosis, and the integrity of the plasma membrane. The data presented here continue to unravel the complex and intricate processes of pH-sensing in a relevant human fungal pathogen. These studies will further elucidate general mechanisms by which cells respond to and internalize extracellular stress signals.

**Author Summary:** The work described here explores the genetics and mechanics of a cellular signaling pathway in a relevant human fungal pathogen, *Cryptococcus neoformans*. The findings presented in this manuscript untangle the complex interactions involved in the activation of a fungal-specific alkaline response pathway, the Rim pathway. Specifically, we find that *C. neoformans* is able to sense an increase in pH within the human host, internalize a membrane-bound pH-sensor, and activate a downstream signaling pathway enabling this pathogen to adapt to a novel host environment and effectively cause disease. Revealing the mechanisms of Rim pathway activation within the larger context of the fungal cell allows us to understand how and when this microorganism interprets relevant host signals. Furthermore, understanding how this pathogenic organism converts extracellular stress signals into an adaptive cellular response will elucidate more general mechanisms of microbial environmental sensing and stress response.

## Introduction

The ability for organisms to effectively recognize and transmit signals relating to changes in the external environment is essential for their survival. For microscopic fungal organisms, the ability to specifically sense increases in extracellular pH is known to be important for the production of secondary metabolites [1], the maintenance of the fungal cell wall [2–6], and virulence in the case of fungal pathogens [7–12]. In many fungi, pH recognition processes include the fungal-specific Rim/Pal alkaline response pathway, [7,8,11,13,14]. In the context of this signaling pathway, extracellular pH signals are initiated through cell surface pH-sensing complexes, which include the Rra/Rim/Pal putative sensors. These signals are then transduced through Endosomal Sorting Complex Required for Transport (ESCRT)-dependent trafficking. Further processing of these alkaline signals is completed through the formation of a proteolysis complex required for cleavage and activation of the Rim101/PacC transcription factor, the terminal component of the pathway [11]. This protein in turn controls the transcriptional activation of numerous genes directing pH-mediated adaptive responses.

Many of the components of the Rim/Pal pathways are highly conserved across diverse fungal phyla including the involvement of the ESCRT machinery and formation of the proteolysis complex. However, the specific pH-sensing proteins present at the cell surface appear to have diverged in a phylum-dependent manner. For example, fungi in the Ascomycota phylum possess plasma membrane-associated pH-sensing proteins with a high degree of sequence and structural similarity – the *Saccharomyces cerevisiae* and *C. albicans* Rim21 proteins, and the orthologous *Aspergillus fumigatus* and *A. nidulans* PalH proteins. Each of these contain seven membrane-spanning domains and a cytoplasmic C-terminal domain [8,11].

The opportunistic fungal pathogen *C. neoformans* is a notable cause of lethal infections in highly immunocompromised patients, especially those with advanced HIV disease [15]. In contrast to many other fungal pathogens of humans, *C. neoformans* belongs to the phylum Basidiomycota, along with many agricultural pathogens and mushrooms. Rim21 homologs are conspicuously absent from the genomes of the basidiomycete fungi [8]. We recently identified the *C. neoformans* Rra1 protein as the most upstream component of the *C. neoformans* Rim pathway, likely serving as the surface alkaline pH sensor [8]. Even though it possesses no sequence similarity to Rim21, *C. neoformans* Rra1 is also predicted to contain seven transmembrane domains and a cytoplasmic C-terminal tail, suggesting functional similarity. Also like Rim21 proteins, Rra1 localizes to the plasma membrane in punctate structures during growth at low pH [16]. At the plasma membrane, this pH sensor is stabilized by the Nucleosome Assembly Protein 1 (Nap1) chaperone [17]. When exposed to alkaline growth conditions, Rra1 senses a pH-induced shift in phospholipid distribution and charge within the plasma membrane, allowing for its highly charged C-terminal tail to disassociate from the inner leaflet into the cytosol [16]. A similar model of plasma membrane-induced activation of the *S. cerevisiae* Rim21 pH sensor has also been suggested [18]. The structural and functional similarities between these highly diverged pH-sensing proteins suggests convergent evolution of the most proximal components of fungal pH-sensing between divergent fungal phyla.

The formation of Rra1 membrane-associated puncta at low pH initially led us to further investigate the connection between Rim pathway activation and plasma membrane dynamics. We have previously shown that the disruption of lipid rafts in the membrane results in mislocalization of the Rra1 pH sensor and hypothesized that Rra1 membrane localization is connected to the formation of distinct membrane domains [16]. While the connection between extracellular stress and membrane dynamics has been made in *C. neoformans* [19–22], these associations were the first to connect the Rim pathway and the plasma membrane in this fungal pathogen. Furthermore, they have revealed potential connections between Rra1 receptor cycling and pH sensing in general fungal virulence.

Several questions remain unanswered regarding microbial/fungal sensing of extracellular pH. These include how fungal plasma membrane pH sensors, like *C. neoformans* Rra1, become internalized in response to changes in environmental pH. Also, it is not yet known how changes in Rra1 protein localization affect Rra1 function and Rim pathway activation. Here we show that *C. neoformans* Rra1 undergoes endocytosis following a shift to alkaline growth conditions and that this endomembrane localization is important for Rim pathway activation. We observe that inhibiting the ability of Rra1 to aggregate at the plasma membrane in acidic conditions does not affect downstream Rim pathway function or growth at alkaline pH. Furthermore, through protein interaction studies, inhibition experiments, and genetic epistasis, we find that this internalization mechanism involves clathrin-mediated endocytosis and phosphorylation of the Rra1 C-terminal tail. Finally, detailed phospholipidomics studies connect the Rim-mediated pH response with the content of cellular membranes. The studies presented here continue to inform the intricate mechanism by which this human fungal pathogen senses and responds to changes in its environment, specifically that of the relatively alkaline human host.

## Results

### Rra1 is endocytosed in response to alkaline pH and recycled back to the membrane

Our previous studies identified Rra1 as a membrane-associated upstream component of the Rim alkaline response pathway in *C. neoformans*. Specifically, we observed that Rra1 is required for Rim pathway activation and growth at alkaline pH [8] and has a pH-dependent localization pattern [16]. Furthermore, the Rra1 C-terminal cytoplasmic tail plays an important role in the localization and function of this putative pH-sensing protein by its differential affinity with the plasma membrane at different pH’s [16].

In order to better define *C. neoformans* Rra1 pH-dependent localization, we examined a detailed time course of Rra1 trafficking in response to alkaline extracellular signals. We used FM4-64, a dye that tracks endocytic transport from the plasma membrane, and assessed the colocalization of this dye with a functional, C-terminally tagged Rra1-GFP fusion protein (Rra1-GFP) [16,23]. In acidic conditions (non-Rim activating conditions), we observed Rra1 enriched in puncta at the cell surface [16]. A shift from pH 4 to pH 8 resulted in reproducible patterns of pH-dependent changes in Rra1 localization. After 10 minutes of exposure to alkaline pH, Rra1-GFP begins to migrate from its sites of plasma membrane aggregation to internal cytoplasmic structures (Fig 1A). The specific foci of Rra1 internalization colocalize with the FM4-64 dye, suggestive of endocytic vesicles (Fig 1A). After extended incubation (20 minutes) at alkaline pH, Rra1 localization changes from surface-associated puncta to endomembranes, including a perinuclear enrichment consistent with the perinuclear endoplasmic reticulum (ER) (Fig 1A). Following endocytosis, FM4-64 follows similar patterns of colocalization with Rra1 on these endomembrane structures (Fig 1A). Furthermore, following activation and endocytosis, we observed that Rra1 recycles back to the cell surface. Specifically, when cells are incubated in alkaline conditions (pH 8) and then re-exposed to pH 4 growth conditions, Rra1 repositions itself in plasma membrane-associated puncta similarly to the original localization pattern observed at pH 4 (Fig 1B and 1C). We also observed that this recycling efficiency is significantly decreased in the *rim101*Δ mutant strain. In the absence of Rim101, there is a delay in the reestablishment of Rra1 enrichment in cell surface puncta following a shift from alkaline to acidic pH (Fig 1B and 1C). Overall, this data revealed that Rra1-GFP undergoes endocytosis from the cell surface to endomembranes in response to alkaline pH and that this protein recycles back to the cell surface following activation.

**Fig 1.**
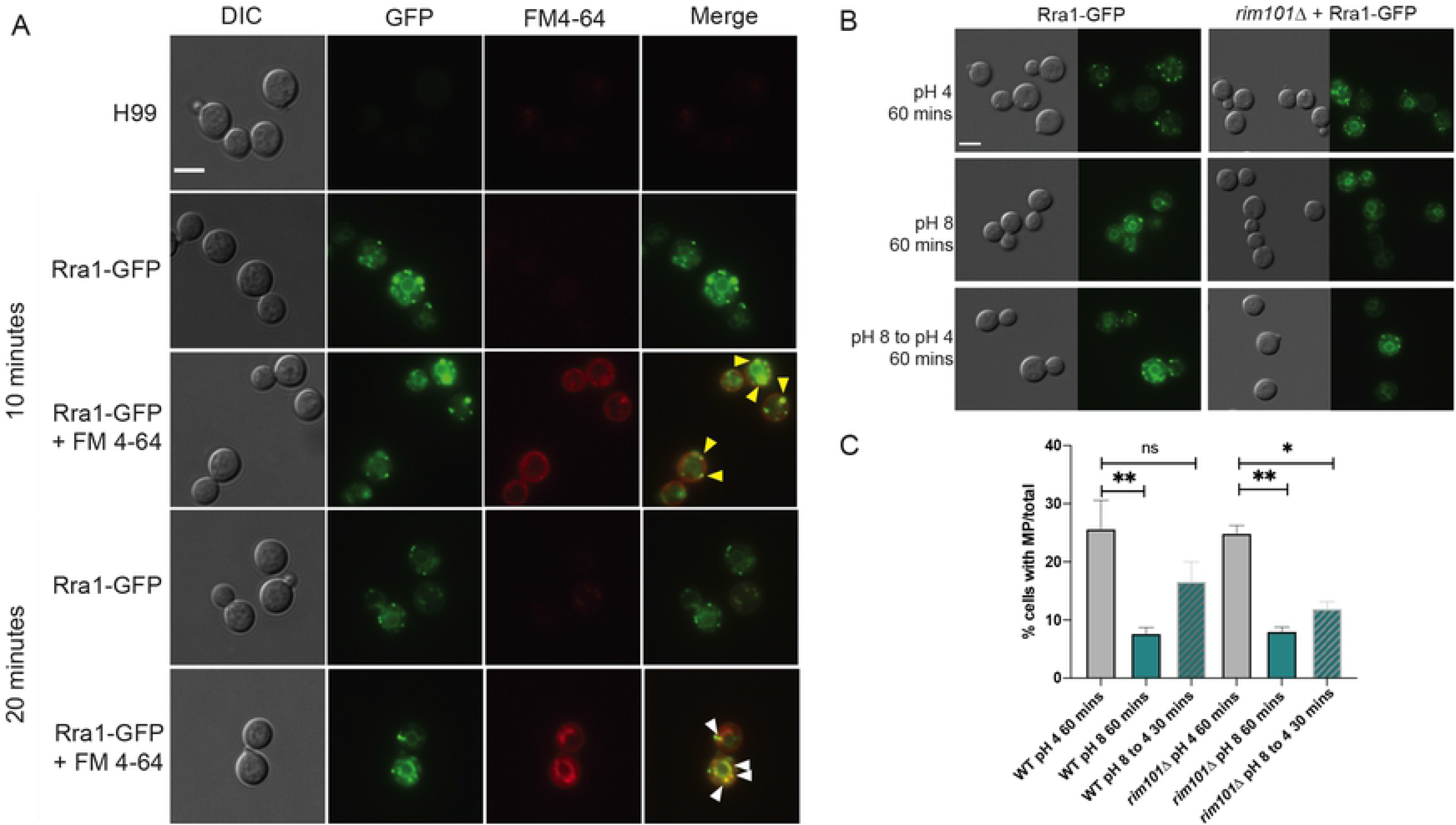
Rra1 colocalizes with FM4-64 labeled structures. A. The wildtype strain with GFP labeled Rra1 was treated with FM4-64 dye following a shift from pH 4 to pH 8 (SC medium buffered to pH 4 and 8 with Mcllvaine’s buffer, referred to as McIlvaine’s medium) at room temperature. Localization of Rra1 (green) and FM4-64 (red) was visualized using epifluorescence microscopy at 10 and 20 minutes. Rra1-GFP colocalization events with FM4-64 near the plasma membrane are indicated by yellow triangles. Colocalization on endomembrane structures is indicated by white triangles. White scale bars indicate 5 microns. B. pH-dependent localization and recycling of the Rra1-GFP fusion construct. The Rra1-GFP strain was incubated at pH 4 or pH 8 McIlvaine’s media for 60 minutes and then shifted back to pH 4 media for 30 minutes in the wildtype and *rim101*Δ strains. GFP signal was assessed by epifluorescence microscopy (Zeiss Axio Imager A1) using the appropriate filter. White scale bars indicate 5 microns. C. Quantification of Rra1-GFP cell surface puncta at pH 4 and pH 8. The mean values and standard errors of cells with > 2 membrane puncta (MP) formed at pH 4 and 8 McIlvaine’s media for 60 minutes and then shifted back to pH 4 for 30 minutes was quantified using ImageJ software (Fiji) (~600 cells/condition; 3 biological replicates). One-way ANOVA, Tukey’s multiple comparison test: ** = p = 0.0014, * = p = 0.0267, ns = not significant.

### Rra1 pH-dependent endocytosis is clathrin-dependent

We assessed the effect of the Pitstop-2 clathrin-mediated endocytosis (CME) inhibitor on Rra1 pH-induced endocytosis. Cells expressing Rra1-GFP were treated with either Pitstop-2 or DMSO vehicle control in pH 4 and pH 8 growth conditions. Following a 10-minute Pitstop-2 treatment, we observed alterations in the endocytosis of Rra1 at pH 8. We noted accumulation of Rra1 in globular structures near the plasma membrane as well as a lack of expected alkaline pH-mediated endomembrane localization (Fig 2A and 2B). These results indicate that Pitstop-2 clathrin inhibition disrupts alkaline pH-induced perinuclear ER localization of the Rra1 protein. In contrast, CME inhibition with Pitstop-2 did not lead to a significant alteration in membrane puncta at pH 4 (Fig S1).

**Fig 2.**
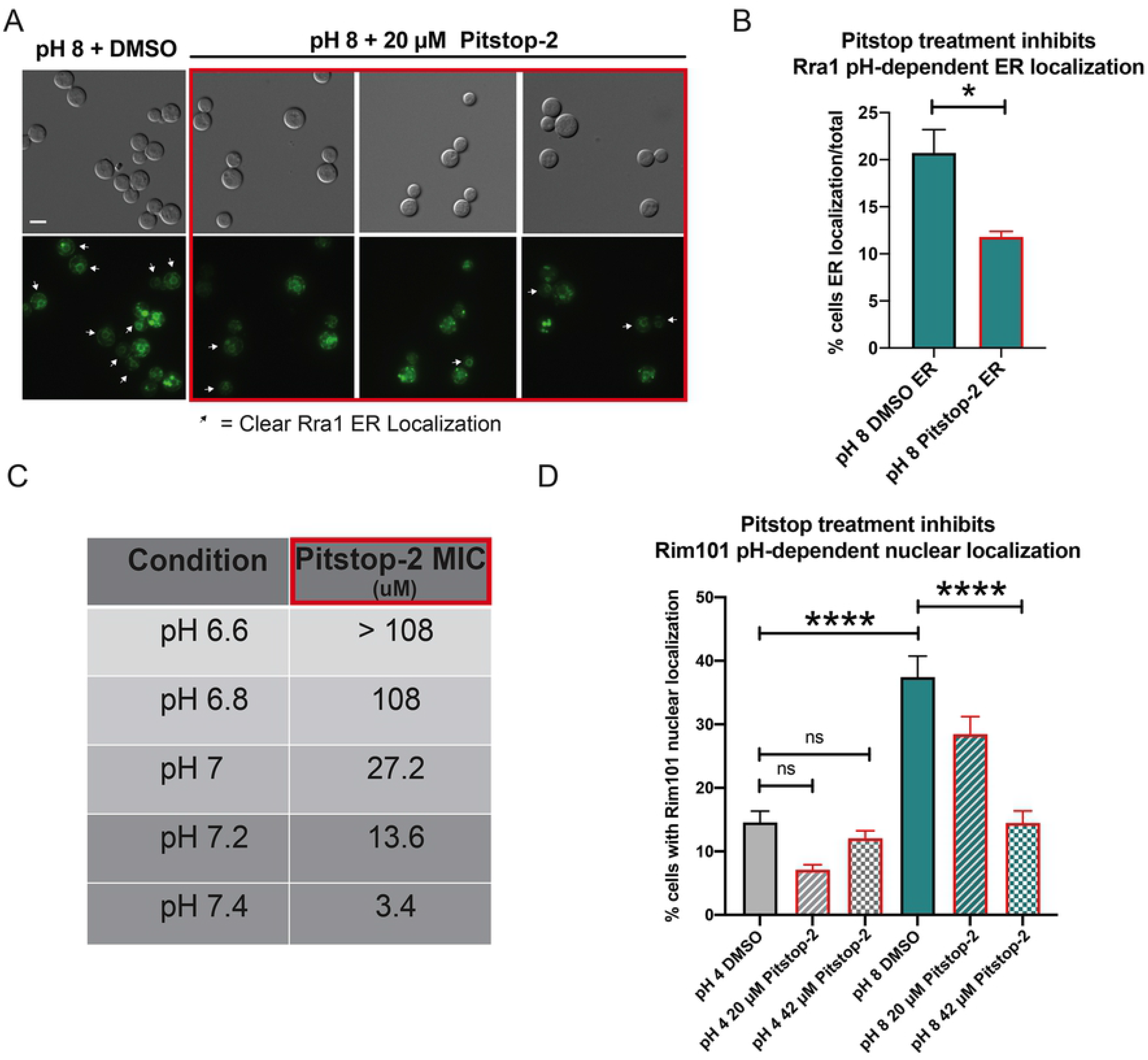
Pitstop-2 inhibition of CME affects Rim signaling. A. Alterations in pH-dependent localization of the Rra1 protein GFP fusion protein by inhibition of CME in response to pH 8 McIlvaine’s medium for 10 minutes following treatment with either 20 μM Pitstop-2 or DMSO. GFP signal was assessed by epifluorescence microscopy (Zeiss Axio Imager A1) using the appropriate filter. White arrows indicate clear endomembrane/ER localization. White scale bars indicate 5 microns. B. Quantification of Rra1-GFP localization in pH 8 McIlvaine’s medium. The mean values and standard errors of cells with clear ER localization at pH 8 was quantified using ImageJ software (Fiji) (~600 cells/condition; 4 biological replicates). Student’s *t*-test, p = 0.012. C. Assessment of MIC of Pitstop-2 CME inhibitor on wildtype *C. neoformans* cells grown under increasingly alkaline conditions. MIC was determined after 72 hours of growth at 30°C by broth microdilution. D. Quantification of pH-dependent nuclear localization of the Rim101 transcription factor in response to pH 4 and pH 8 McIlvaine’s media for 10 minutes following treatment with either Pitstop-2 (20 μM or 42 μM) or DMSO. The mean values and standard errors of cells with clear nuclear localization at pH 8 was quantified using ImageJ software (Fiji) (~600 cells/condition; 3 biological replicates). One-way ANOVA, Tukey’s multiple comparison test: **** = p < 0.0001. White scale bars indicate 5 microns.

To assess whether Pitstop-2 treatment and its associated alterations in Rra1 localization affect growth at alkaline pH, we incubated wildtype *C. neoformans* cells at a range of pH levels and exposed to increasing concentrations of Pitstop-2 for 48 hours. In addition to the associated changes in Rra1 localization, clathrin inhibition with Pitstop-2 also resulted in functional consequences for growth at elevated pH. Low concentrations of Pitstop-2 (3.4 μM) inhibited fungal growth at an alkaline pH (YPD pH 7.4). However, *C. neoformans* was able to grow at much higher concentrations of this clathrin inhibitor (> 108 μM) in a slightly more acidic medium (YPD pH 6.6) (Fig 2C).

In order to directly assess whether blocking CME leads to defective Rim pathway signaling, we tested the effects of Pitstop-2 on the nuclear translocation of the Rim101 transcription factor in response to increases in pH. Rim101 is the terminal transcription factor in the Rim pathway, and its translocation to the nucleus following a shift to alkaline pH is a hallmark of pathway activation [8]. We observed a dose-dependent decrease in pH-regulated Rim101 nuclear localization following Pitstop-2 treatment compared to vehicle treated cells (Fig 2D). Together these data indicate that blocking CME results in alkaline pH sensitivity, likely through inhibition of both Rra1 endocytosis and subsequent Rim101 nuclear translocation.

### Rim pathway upstream components interact with endocytosis machinery at alkaline pH

To further assess Rra1 trafficking and interactions of this protein with downstream effectors, we performed mass spectrometry on proteins co-immunoprecipitated with the Rra1 C-terminus. The Rra1 C-terminus is a soluble subdomain of the Rra1 protein that we have previously shown to be required for Rim signal initiation [16]. Focusing on interactors of this domain avoids the need for strong membrane protein-extracting detergents that might be required for isolation of membrane proteins, but that also might disrupt physiologically relevant protein interactions. We were most interested in proteins that interact with the Rra1 C-terminus in Rim pathway-activating conditions (alkaline pH); therefore, we performed a co-immunoprecipitation using a GFP-tagged version of the Rra1 C-terminus (GFP-Rra1-Ct) at pH 8. The GFP-Rra1-Ct was immunoprecipitated from cell lysates using a GFP-Trap resin, and the associated proteins were analyzed using tandem MS-MS. To exclude potential false-positive interactions, we prioritized proteins with at least 5 exclusive peptides that were present only in the GFP-Rra1-Ct sample and not in the control condition (Table S1). At pH 8, Rra1 C-terminus interactors included proteins typically found on endocytic vesicles, such as coatomer protein subunits, clathrin heavy chain 1, and archain 1 (Table 1). Also included were multiple T-complex protein subunits that are typically found to interact with endomembrane-associated proteins (*i.e*., secretory proteins (Sec27) and COP proteins). The Nap1 chaperone protein was also found to be a strong interactor with the Rra1 C-terminus at high pH, supporting our previous studies revealing that Nap1 stabilizes the Rra1 protein, specifically through its interaction with the C-terminus [17]. Furthermore, gene ontology analysis using FungiFun FunCat [24], revealed protein fate (i.e. protein folding, modification, and destination) as one of three categories significantly represented in the Rra1-Ct interactome (blue font in Table 1 and Fig 3A), and COPI-vesicle coat as one of the significant cellular compartment GO-term categories (red font in Table 1 and Fig 3B). These results are consistent with our findings outlined above regarding the clathrin-mediated endocytic trafficking of Rra1 to endomembrane sites of downstream activity (Fig 2). Furthermore, a previously published protein interaction study assessing proteins co-immunoprecipitated with the full-length Rra1-GFP in alkaline conditions identified COPI and clathrin subunits among the interacting partners (Table S1 in [17]). This supports the role for endocytosis machinery in the internalization of Rra1 in alkaline conditions.

**Fig 3.**
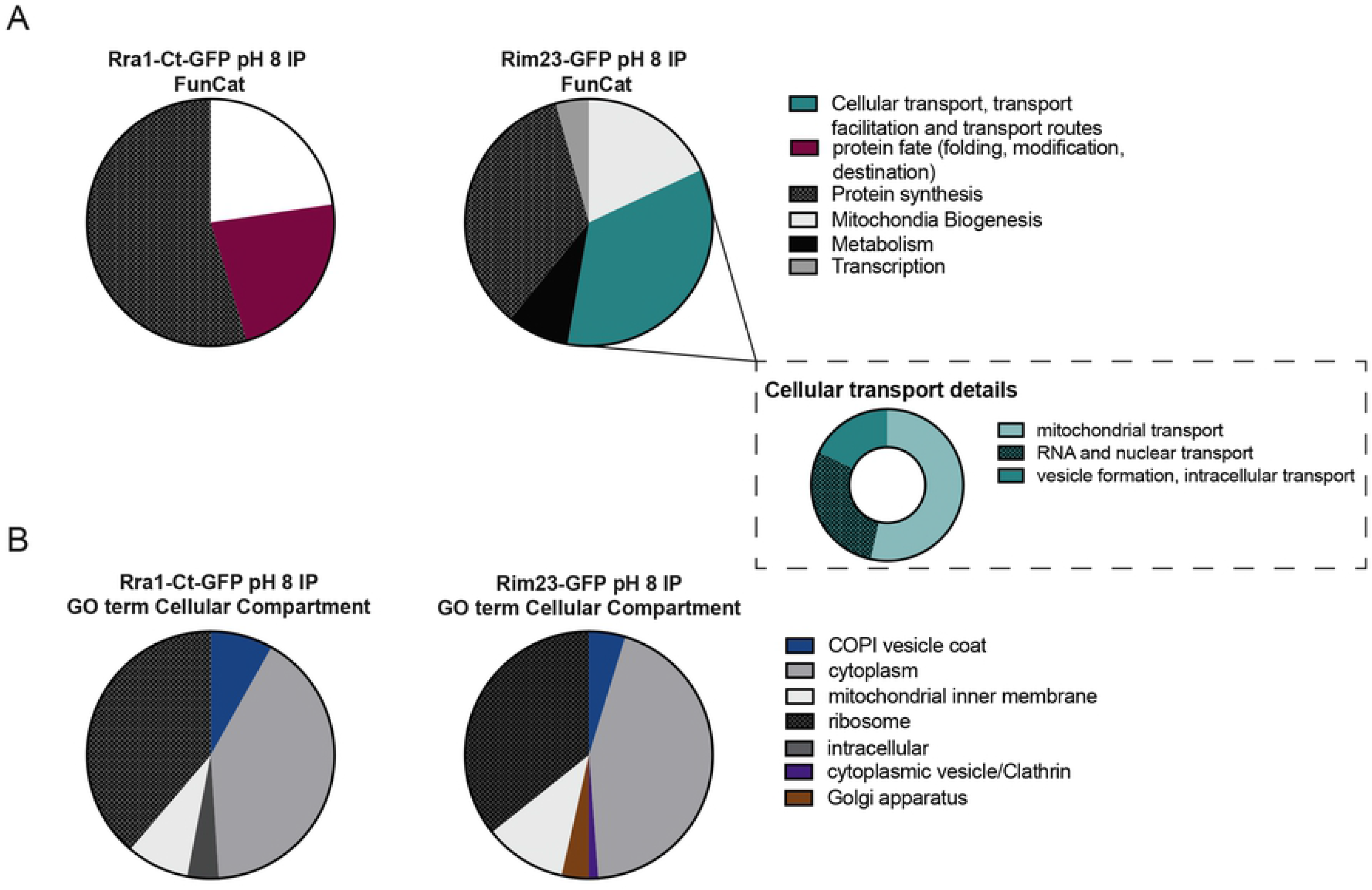
Upstream Rim pathway components interact with endocytosis machinery at high pH. Following incubation of the GFP-Rra1-Ct and the Rim23-GFP expressing strains in alkaline conditions (YPD pH 8) for one hour, cell lysates were immunoprecipitated using a GFP-Trap resin. The associated proteins were analyzed using tandem MS-MS. These interactomes were then analyzed with FungiFun software to identify significantly enriched Gene Ontology categories. (A) FunCat analysis from the Rra1-Ct and Rim23 interactomes and the inset of the Rim23 FunCat results represents the subcategories within the umbrella cellular transport. (B) GO-term analysis on the enriched cellular compartments for the two interactomes. The specific CNAG #s and gene names in each category can be found in Tables 1 and 2, and the full interactomes can be found in Table S1.

**Table 1.** Proteins enriched in Gfp-Rra1-Ct interactome at pH8 compared to untagged control.

| Gene ID | Gene name/predicted ortholog | Peptide counts |
| --- | --- | --- |
| CNAG_01274 | Coatomer subunit gamma | 8 |
| CNAG_03299 | Coatomer subunit beta | 13 |
| CNAG_03554 | Coatomer subunit alpha | 6 |
| CNAG_02937 | Hypothetical protein | 5 |
| CNAG_04074 | Coatomer subunit beta' | 28 |
| CNAG_01148 | Peptidyl prolyl isomerase | 6 |
| CNAG_00058 | T-complex protein subunit | 7 |
| CNAG_00447 | T-complex protein subunit | 6 |
| CNAG_01019 | Cu superoxide dismutase | 5 |
| CNAG_01568 | T-complex protein subunit | 10 |
| CNAG_02038 | Hypothetical protein | 9 |
| CNAG_02710 | T-complex protein subunit | 14 |
| CNAG_03459 | T-complex protein subunit | 12 |
| CNAG_04304 | T-complex protein subunit | 21 |
| CNAG_05105 | translation initiation factor | 7 |
| CNAG_06600 | Vesicular protein Sec18 | 6 |
| CNAG_07347 | Heat shock protein | 9 |
| CNAG_06508 | Glucan synthase Fks1 | 6 |
| CNAG_02091 | Nucleosome protein Nap1 | 19 |
| CNAG_04613 | ATP-dependent transporter | 13 |
| CNAG_02753 | ER protein | 6 |
| CNAG_06630 | Membrane translocase Tim44 | 5 |

This subset of potential Rra1-Ct interactors is involved in intracellular protein transport. GO-term cellular component, FunCat, both.

As mentioned previously, Rim signaling is initiated by the formation of a Rra1-containing cell surface pH-sensing complex, and it is completed through the formation of a proteolysis complex required for Rim101 cleavage. Rim23 is a component of the proteolysis complex, and this protein displays membrane-associated localization in response to a shift to alkaline pH [8]. Therefore we were also interested in the interactome of this protein in activating conditions and whether the Rra1-Ct and Rim23 complexes might interact. We performed a similar protein interaction study with a GFP-tagged version of Rim23 in alkaline conditions. Similar to those with GFP-Rra1-Ct, Rim23 interactors were also enriched for coatomer and clathrin-associated proteins at pH 8 (Table 2). FungiFun FunCat gene ontology analysis of the Rim23 interactome revealed cellular transport, transport facilitation, and transport routes as significantly enriched categories [24]. These proteins included those involved in vesicle formation and intracellular transport such as coatomer subunits, clathrin protein Ap47, clathrin heavy chain, and transport protein Sec13 (blue font Table 2 and Fig 3B). Additionally, the significantly enriched cellular component GO term categories consisted of COPI-vesicle coat, cytoplasmic vesicle, clathrin, and Golgi apparatus (red font Table 2 and Fig 3D). These results indicate that the Rim Sensing/Activation Complex and the Rim Proteolysis Complex likely physically and temporally converge at common sites during pathway activation and that these sites contain proteins involved in protein trafficking and CME. We have not observed a similar pattern of enrichment of endocytic vesicle-associated proteins in other proteomics experiments [25].

**Table 2.** Proteins enriched in Rim23-Gfp interactome at pH8 compared to untagged control.

| Gene ID | Gene name/predicted ortholog | Peptide counts |
| --- | --- | --- |
| CNAG_01274 | Coatomer subunit gamma | 18 |
| CNAG_03299 | Coatomer subunit beta | 26 |
| CNAG_03554 | Coatomer subunit alpha | 48 |
| CNAG_01414 | ARCN1 protein | 8 |
| CNAG_00988 | Importin subunit beta-1 | 33 |
| CNAG_02457 | Importin subunit beta-2 | 20 |
| CNAG_03317 | Clathrin protein Ap47 | 10 |
| CNAG_03418 | Crm1-F1 | 13 |
| CNAG_03853 | COP11 GTPase Sar1 | 11 |
| CNAG_04074 | Coatomer subunit beta | 20 |
| CNAG_04904 | Clathrin heavy chain 1 | 16 |
| CNAG_05884 | Importin subunit beta-4 | 20 |
| CNAG_06630 | Translocase subunit Tim44 | 13 |
| CNAG_07318 | Gamma-adaptin | 6 |
| CNAG_07598 | Import-alpha receptor | 7 |
| CNAG_01211 | Hypothetical protein | 11 |
| CNAG_04194 | Protein transport Sec13 | 9 |
| CNAG_06600 | Protein transport Sec18 | 30 |
| CNAG_01837 | Vacuolar sorting Vps35 | 13 |
| CNAG_06998 | Protein transporter | 8 |
| CNAG_00539 | Membrane transport | 7 |
| CNAG_01637 | CopII Vesicle Erv46 | 5 |
| CNAG_07570 | Snare protein Ykt6p | 5 |
| CNAG_01426 | Vacuolar sorting Vps26 | 5 |

This subset of potential Rim23 interactors is involved in intracellular protein transport. GO-term cellular component, FunCat, both

### Rra1 pH dependent localization is altered through disruption in membrane composition

We previously identified a Rim-independent mechanism of the fungal alkaline pH response in which the Sre1 transcription factor and its downstream effectors in the ergosterol biosynthesis pathway are activated in response to alkaline pH [26]. Other work has also demonstrated that the *sre1*Δ mutant has depleted levels of ergosterol in the plasma membrane and altered abundance of sterol-rich domains, affecting the localization of membrane-associated proteins [27–29]. We also previously observed that altering the formation of lipid rafts in the membrane using Filipin III dye results in disruption of Rra1 membrane puncta formation at pH 4 [16]. We therefore assessed the effects of Sre1 mutation on the localization of Rra1.

In contrast to wildtype, Rra1 membrane-associated puncta were not observed at pH 4 in the *sre1*Δ mutant strain. In this mutant background, Rra1 is localized to endomembranes in both activating (pH 8) and inactivating conditions (pH 4) (Fig 4A and 4B). However, Rim signaling is still intact in the *sre1*Δ mutant background as demonstrated previously by normal processing of the Rim101 transcription factor in response to elevated pH [26]. Together these data support that Rra1 membrane puncta are not essential for alkaline-induced Rim signaling. Furthermore, treating wildtype cells with Filipin III does not lead to decreased growth at alkaline pH despite similar disruption of cell surface puncta. Wildtype *C. neoformans* cells were able to grow at a range of increasing pH growth conditions (pH 4,5,6,7, and 8) despite high concentrations of Filipin III (62.5 ug/mL) [10 ug/mL for microscopy experiments in [26]]. These results indicate that Sre1-mediated ergosterol and membrane homeostasis is essential for Rra1 localization in plasma membrane puncta at low pH, but that this localization is not necessary for Rim pathway activation.

**Fig 4.**
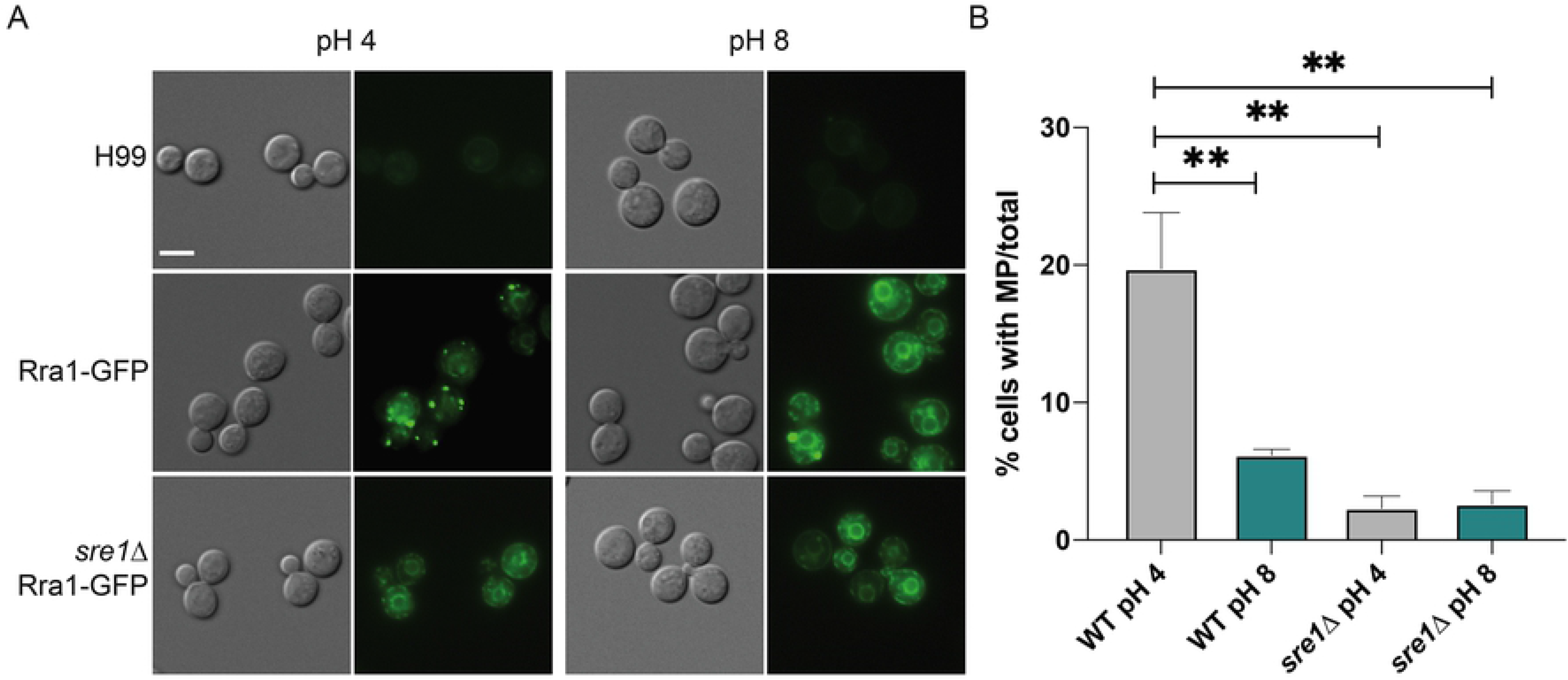
Reduced Rra1-containing membrane puncta at low pH in the *sre1*Δ mutant strain. A. pH-dependent localization of the Rra1-GFP protein fusion construct in response to pH 4 and pH 8 McIlvaine’s media for 60 minutes in the wildtype and *sre1*Δ mutant backgrounds. GFP signal was assessed by epifluorescence microscopy (Zeiss Axio Imager A1) using the appropriate filter. White scale bars indicate 5 microns. B. Quantification of Rra1-GFP localization at pH 4 and pH 8. The mean values and standard errors of cells with > 2 membrane puncta formed at pH 4 and 8 was quantified using ImageJ software (Fiji) (~600 cells/condition; 3 biological replicates). One-way ANOVA, Tukey’s multiple comparison test: ** = p < 0.0095.

### Assessment of Rra1 C-terminus pH-dependent structure and phosphorylation

Our recently published studies suggest that the C-terminal tail of Rra1 serves as an “antenna” to mediate pH-dependent interactions with the plasma membrane [16]. These results are further supported through Rra1 structural predictions using various modeling platforms. Two major structural models emerge from the amino acid sequence of the Rra1 protein: one that maintains the C-terminal region tightly compact and one that displays a free and extended C-terminus (Fig S2) [30–32]. These two orientations of the Rra1 C-terminus might represent the bimodal function of this domain as it differentially interacts with the plasma membrane in response to changes in charge of the inner leaflet [16,18]. Furthermore, protein truncation studies demonstrated that the Rra1 C-terminus, and especially the highly charged region (HCR), as graphically represented in Fig 5A, is required for the function of this protein. A mutated form of Rra1-GFP lacking the entire C-terminus after residue 273 [Rra1-273T-GFP (T = truncated)] was unable to restore alkaline growth to the *rra1*Δ mutant (Fig 5F and [16]. In contrast, a truncated Rra1-GFP protein that retained the HCR (Rra1-296T-GFP) completely complemented *rra1*Δ mutant phenotypes (Fig 5F and [16]. This truncated strain revealed localization patterns that mirrored wildtype, however we later learned that this strain also contained a full-length *RRA1-GFP* allele. Repeating these localization in a new strain, with the truncated Rra1-296T-GFP as the cellular source of Rra1, revealed similar localization patterns to wildtype with Rra1-containing membrane puncta at low pH and Rra1 internalization at high pH, identical to the previously published results [16]. Interestingly, we noted that at high pH, this truncated strain appeared to have increased levels of Rra1 in endomembrane structures consistent with the robust growth of this strain at high pH (Fig 5F).

**Fig 5.**
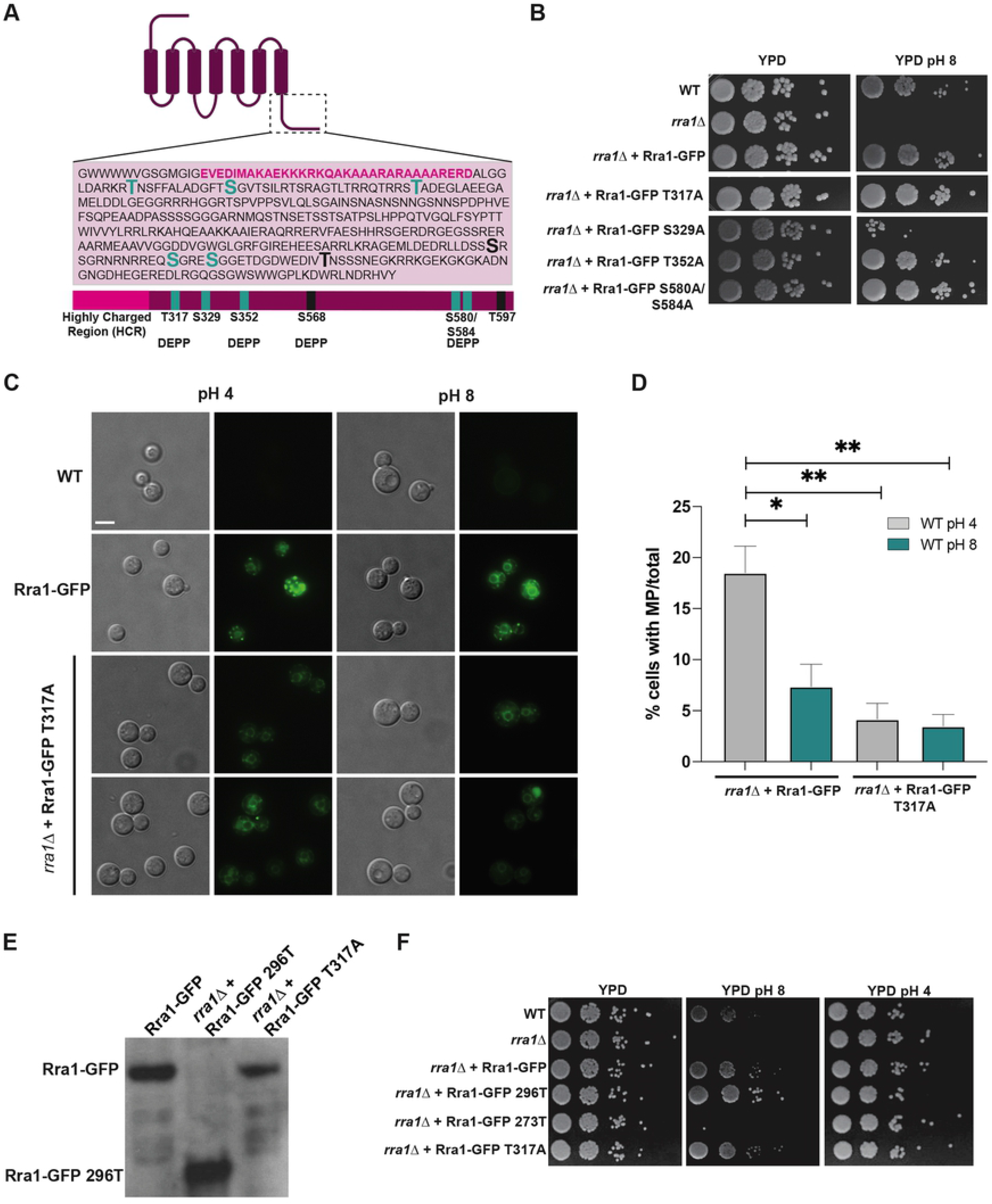
Rra1 phosphomutant affects Rra1 localization, but not function. A. Schematic of the pH-dependent phosphosites of the Rra1 protein. Sites that are preferentially phosphorylated at pH 4 are depicted in black and sites preferentially phosphorylated at pH 8 are depicted in teal. Sites that were also predicted to be phosphorylated using DEPP software are labeled with “DEPP”. The highly charged region that has been shown to be essential for Rra1 function and proper localization is indicated in fuchsia. B. The prioritized alkaline phosphorylation site mutants, incubated overnight in YPD medium, were serially diluted onto YPD and YPD pH 8 agar plates to assess growth rate compared to wildtype, the *rra1*Δ mutant strain, and the Rra1-GFP strain. Plates were incubated at 30° C for 3 days prior to imaging. C. The Rra1-GFP wildtype and the Rra1-GFP T317A phosphomutant strains were incubated in pH 4 and pH 8 McIlvaines media for 60 minutes. Rra1-GFP localization was assessed by epifluorescence microscopy (Zeiss Axio Imager A1) using the appropriate filter. White scale bars indicate 5 microns. D. Quantification of Rra1-GFP localization at pH 4 and pH 8 in the Rra1-GFP wildtype and T317A phosphomutant backgrounds. The mean values and standard errors of cells with > 2 membrane puncta formed at pH 4 (grey) and 8 (teal) McIlvaine’s buffer for 60 minutes was quantified using ImageJ software (Fiji) (~600 cells/condition; 3 biological replicates). One-way ANOVA, Tukey’s multiple comparison test: * = p = 0.0165, ** = p < 0.0038 E. Western blot analysis of Rra1 protein levels in different genetic backgrounds: wildtype, the Rra1-296T truncation mutant that retains the HCR, and the T317A phosphomutant. Strains were incubated for 1.5 h in pH 8 YPD buffered with 150 mM HEPES. Samples were assessed by western blotting using an α-GFP antibody. White scale bars indicate 5 microns. F. Comparison of the Rra1-GFP T317A phosphomutant to the truncation mutants, *rra1*Δ and wildtype strains and their respective growth on acidic and alkaline pH. Strains were serially diluted onto YPD and YPD pH 8, and YPD pH 4 agar plates to assess growth rates in pH stress. Plates were incubated at 30° C for 3 days prior to imaging.

Given the central role for the exposed Rra1 C-terminus in protein function, we hypothesized that pH-dependent post-translational modifications (PTMs) of the Rra1 protein, specifically within the C-terminus, would direct its localization and function. We therefore assessed Rra1 phosphorylation patterns at two extremes of pH: pH 4 (Rim pathway non-activating) and pH 8 (Rim pathway activating). We chose to focus on this specific PTM based on (1) DEPP and PONDR prediction software revealing the Rra1 C-terminus to be highly disordered and positioned for phosphorylation modifications (Fig 5A and Fig S2) [33,34] (2) our identification of this region of the Rra1 protein as the site of interaction with downstream proteins such as Nap1 ([17] and Table 1) and (3) preliminary MS analysis demonstrating pH-dependent changes in Rra1 phosphorylation as described in our methods.

As graphically depicted in Fig 5A, we observed two different patterns of Rra1 protein serine/threonine phosphorylation: residues preferentially phosphorylated at pH 8 and residues phosphorylated at pH 4. Interestingly, all pH-dependent changes in Rra1 phosphorylation were present in the cytoplasmic C-terminal tail (Fig 5A). To assess the role of each potential phosphosite on Rim-regulated cellular functions, we created *RRA1* alleles with alanine mutations at each of these serine or threonine residues. We prioritized strains with alanine substitutions in residues preferentially phosphorylated at alkaline pH (Fig 5B). For each strain, we assessed fluorescent protein localization (epifluorescence microscopy), transcript and protein stability (RT-PCR and western blots, respectively), and complementation of *rra1*Δ growth defects at pH 8 (Fig 5B). Most of these mutations did not alter Rra1-GFP localization or function. The one phosphomutant that did affect the ability to grow at alkaline pH (Rra1-GFP-S329A) displayed unstable *RRA1* transcript levels at pH 8 and therefore was not prioritized. However, in contrast to the wildtype Rra1-GFP that localized in PM puncta at acidic pH, one phosphomutant strain (Rra1-GFP-T317A) displayed reduced plasma membrane puncta at low pH, similar to Rra1 localization in the *sre1*Δ mutant (Fig 5C and 5D). We confirmed wildtype expression levels of this mutated protein by western blot (Fig 5E) and wildtype transcript levels by quantitative real-time PCR (Fig S3B). Given its absent Rra1 puncta at low pH and the inability for Rra1 to cycle to and from the PM puncta (Fig S3A), we first hypothesized that this strain would display defective Rim signaling. However, Rra1-T317A fully supported Rim pathway activation as inferred by restoration of growth at alkaline pH as well as acidic pH (Fig 5B and 5F). This intact signaling is similar to the previously published strain lacking the region of the Rra1 C-terminus following the HCR (296T truncation) which involves the removal of the T317 residue (Fig 5B and 5F). Furthermore, this phosphomutant strain displayed a restoration of the alkaline-induced transcriptional induction of *CIG1* expression, which is impaired in Rim pathway mutants (Fig S3C [16]). Together these results strongly suggest that pH-dependent phosphorylation events mediate Rra1 protein localization. They also further support that plasma membrane microdomains, or membrane puncta, are not the sites of Rra1 interaction with its downstream effectors.

### pH-dependent phospholipid analysis

In order to further investigate the effect of membrane composition on pH signaling, we assessed the phospholipid profile of the wildtype strain in response to changes in pH. We hypothesized that if Rra1 cycling through membrane invagination and endocytosis was important for pathway activation and growth at alkaline pH, then the membranes associated with this protein must be changing in a pH-dependent manner to facilitate internalization. This analysis revealed reproducible increases in two out of the five most abundant phosphatidylethanolamine (PE) species in alkaline pH (Fig 6A), and a decrease in 6/13 most abundant phosphatidylserine (PS) and 6/23 most abundant phosphatidylcholine (PC) species in the same alkaline conditions (Fig 6B and 6C, respectively). A majority (10/13) of the most abundant species that were found to be significantly altered in response to alkaline conditions were unsaturated lipids (Fig 6A–6E, indicated by #). Unsaturated phospholipids can sterically hinder the formation of lipid rafts in the plasma membrane.

**Fig 6.**
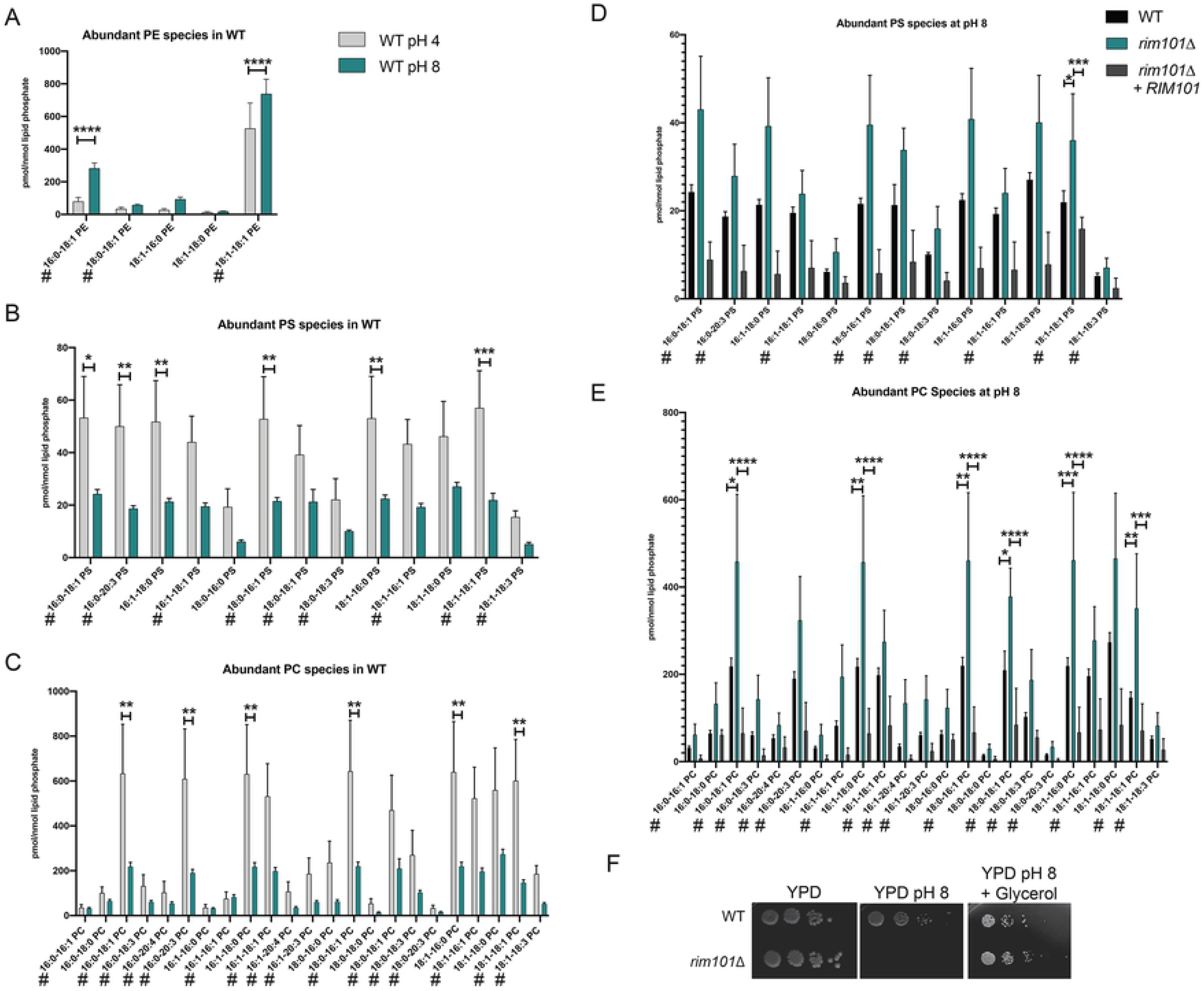
pH-dependent phospholipid analysis. The wildtype *C. neoformans* strain was incubated in pH 4 (grey) or pH 8 (teal) YNB media prior to lipid extraction. Graphs represent lipid profile comparisons of the most abundant (A) phosphatidylethanolamine (PE) (B) phosphatidylserine (PS) and (C) phosphatidylcholine (PC) species analyzed. Two-way ANOVA, Sidak’s multiple comparison test: **** p < 0.0001, *** p < 0.005, ** p < 0.007, * p = 0.01. Statistical tests were run on all lipid species analyzed in biological triplicate using GraphPad Prism. # represents unsaturated lipid species. The wildtype (black), *rim101*Δ mutant (teal) and *rim101*Δ + *RIM101* (dark grey) reconstituted strains were incubated in pH 8 YNB media prior to lipid extraction. Graphs represent lipid profile comparisons of the most abundant (D) phosphatidylserine (PS) and (E) phosphatidylcholine (PC) species analyzed. Two-way ANOVA, Tukey’s multiple comparison test: **** p < 0.0001, *** p < 0.0002, ** p < 0.0021, * p = 0.0332. Statistical tests were run on all lipid species analyzed in biological triplicate using GraphPad Prism. # represents unsaturated lipid species. F) The wildtype and *rim101*Δ mutant strains were serially diluted onto YPD and YPD pH 8 +/- glycerol agar plates to assess growth rate. Plates were incubated at 30° C for 3 days prior to imaging.

Similar phospholipid analysis in the *rim101*Δ mutant revealed increased levels of PC and PS at pH 8 compared to WT and the reconstituted strain. Specifically, at high pH, the *rim101*Δ mutant strain displayed a trend of increased levels of all abundant PC and PS species (Fig 6D and 6E). 5/7 of the statistically significant increases in the most abundant PC and PS species were in unsaturated lipids. These complementary results suggest that the *C. neoformans* Rim pathway is required to maintain pH-induced alterations in the ratios of specific, abundant phospholipids in cellular membranes. These phospholipid alterations are also consistent with previous findings that identified Rim101 as a regulator of the PS decarboxylase (CNAG_00834) in alkaline growth conditions [16]. Furthermore, the *rim101*Δ alkaline pH-sensitive mutant phenotype can be rescued with glycerol supplementation to the growth medium (Fig 6F). Glycerol is the backbone of all phospholipids, and its ability to suppress the severe alkaline growth defect of the *rim101*Δ mutant strain may be due to the re-establishment of normal plasma membrane phospholipid composition. We did not observe a similar trend or any significant differences in PE levels in the *rim101*Δ mutant strain (Table S2).

## Discussion

### Rra1 pH-induced internalization

Endocytosis and protein trafficking from the cell surface allow cells to internalize signals and macromolecules from the extracellular space. Additionally, this process recycles membrane-bound proteins and surrounding lipids [35,36]. Clathrin-mediated endocytosis (CME) is the dominant endocytic pathway in organisms as diverse as mammalian neuronal cells to microbial pathogens. CME has been well-characterized for its role in intracellular communication [37,38], as well as for promoting cellular homeostasis through the internalization of membrane-associated proton pumps and ion channels [37]. CME is initiated by the recruitment of coat proteins and clathrin to membrane-bound receptor-ligand complexes that are targeted for internalization. These coated regions of the membrane invaginate to form endocytic vesicles, which are then transported to intracellular micro-niches including the Endosomal Sorting Complex Required for Transport (ESCRT) (Conner and Schmid, 2003; Gonzá Lez-Gaitá N and Stenmark, 2003; Hurley and Emr, 2006; Miaczynska and Stenmark, 2008; Park *et al*., 2020). Likely due to their involvement in the transport of internalized cellular material, ESCRT proteins are required for stress tolerance, including the adaptation of microbial pathogens to extracellular conditions encountered in the infected human host ([8,23,41–43]. In many fungi such as the human pathogens *C. albicans, A fumigatus*, and *C. neoformans*, and the plant pathogen *Rhizoctonia solani*, this endocytosis process is required for growth and differentiation in response to changes in the extracellular environment [8,23,35,43–45]. Accordingly, disrupting protein trafficking pathways often results in defective fungal virulence [23,43].

Our studies suggest that the *C. neoformans* Rra1 protein is endocytosed in a pH- and clathrin-dependent manner (Fig 7). Furthermore, we have identified this internalization and subsequent enrichment at endomembranes as important for Rim pathway activation. pH-induced endocytosis of transmembrane transporter proteins has been well described in the model ascomycete *S. cerevisiae*. The transporters of inositol (Itr1), uracil (Fur4), tryptophan (Tat2), and hexose (Hxt6) are all endocytosed in response to increases in the bioavailability of their respective substrates. All of these endocytosis events also occur in response to ubiquitination [46]. Endocytosis of Rim-associated proteins has also been explored in other fungi. The *S. cerevisiae* Rim21 protein in *S. cerevisiae* is endocytosed in a pH-dependent manner through a mechanism involving the ubiquitination of the Rim8 arrestin, whose homolog is notably absent from the *C. neoformans* genome [8,47,48]. Furthermore, the Rim21 protein in both *S. cerevisiae* and *C. albicans* colocalizes with ESCRT proteins, such as Snf7, in response to increases in extracellular pH. This interaction between ESCRT proteins and Rim pathway components is required for proteolytic activation of the Rim101 transcription factor [47,49,50].

**Fig 7.**
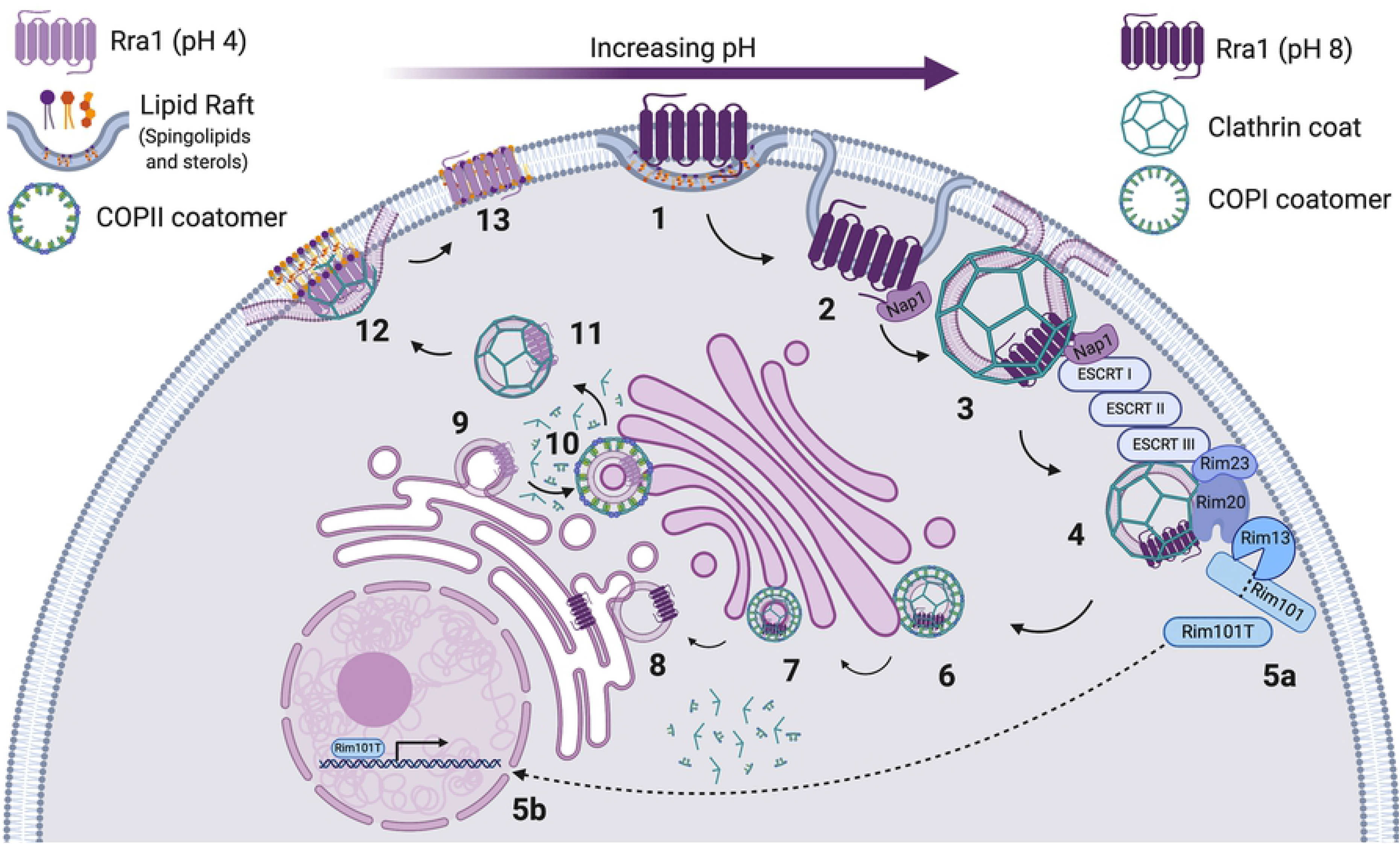
Model of Rra1 cycling resulting in pH-mediated Rim pathway activation. In response to increases in extracellular pH, the Rra1 pH-sensing protein undergoes clathrin-mediated endocytosis from its resting location in sterol-rich PM domains (1-3), re-localizing to endomembranes. At that site, Rra1 assists in the ESCRT-directed assembly of the Rim Proteolysis Complex (4-5a), activating the Rim101 transcription factor to translocate to the nucleus where it controls the expression of its target genes (5b). Recycling back to the PM occurs at more acidic pH (steps 6-13).

pH-dependent endocytosis of pH-sensing proteins is not observed in every fungal organism with a Rim/Pal alkaline response pathway. In *A. nidulans*, studies have definitively shown that Pal signaling and the response to increased pH do not require endocytosis. Using strains with mutations causing varying degrees of endocytosis impairment, the investigators demonstrated intact Rim signaling in these mutant backgrounds [51]. Furthermore, other studies revealed that upstream Pal and ESCRT components in *A. nidulans* localize to cortical plasma membrane puncta in alkaline conditions as opposed to endomembrane structures [52,53]. Therefore, the endocytosis-independent activation of the transmembrane sensor in the *A. nidulans* pathway is distinct from the Rim21 sensor in the *S. cerevisiae* and *C. albicans* pathways. This distinction is especially interesting considering all of these sensors involve ubiquitination-dependent mechanisms of activation via their respective arrestin protein partners. This divergence could be explained by the unique way in which filamentous fungi traffic various proteins. Filamentous fungal membrane transporters can localize in a polar manner, whereas yeast membrane transporters generally localize homogenously in microdomains throughout the plasma membrane [54]. This has been linked to a different path the protein takes following synthesis in the ER. Many of *A. nidulans* transporters bypass the Golgi apparatus and traffic directly to the plasma membrane following synthesis [54]. The ability for proteins to circumvent certain cellular components to a final destination could explain the endocytosis-independent mechanism of Pal pathway activation. The fact the *C. neoformans* Rra1 protein follows similar localization patterns and mechanisms of action as the other yeast-like fungi, but not the filamentous fungi, is compelling considering these proteins lack any sequence homology or evidence of common ancestry [8].

### Rra1 cycles back to the plasma membrane following activation

We have observed that Rra1 returns to the plasma membrane following Rim pathway activation (Fig 7). It is hypothesized that the origin of retrograde sorting, specifically a Golgi-directed pathway originating from the endosome is the key sorting event that allows for plasma membrane recycling of a protein. Our protein interaction studies in the *C. neoformans* Rim pathway in alkaline conditions support a Golgi origin of retrograde sorting for the Rra1 protein (Table 1 and (Ma and Burd, 2020)). These interaction studies also linked the Rra1 C-terminus and the Rim23 protein to clathrin and coatomer proteins in activating conditions.

In yeast, CME-directed internalization of endocytic vesicles is a continuous process, converting half of the material in the plasma membrane to the endosomal system every second [55]. Therefore, cycling of membrane-associated proteins is intimately linked to the plasma membrane. In *S. cerevisiae*, a protein that facilitates vesicle fusion at the cell surface, Snc1, normally recycles from the plasma membrane in a clathrin-dependent manner to the Golgi and then back out through the secretory pathway. However, when depleted from the PM, this protein accumulates in internal organelles [55]. This internal accumulation resembles the Rra1 localization we observed in strains that have been either genetically altered or treated to disrupt plasma membrane composition. This supports our model of Rra1 cycling via clathrin-guided membrane invagination (Fig 7). Additionally, cycling of membrane proteins can be essential for pathway activation. In *S. cerevisiae*, the Cdc42 Rho-GTPase cycles between the membrane and the cytoplasm to regulate cell polarity [56]. This cycling-dependent activation is what we observe with alkaline pH-induced endocytosis of the Rra1 protein and subsequent Rim pathway activation, further supporting our model of Rra1 cycling (Fig 7).

Our results reveal that Rra1 cycling is dependent upon regions of the C-terminal tail and specific phosphorylation events. However, inhibition of this phosphorylation event does not affect growth at alkaline pH or Rim pathway activation despite notable alterations in Rra1 protein localization. PTMs of membrane-associated proteins and their effects on endocytosis and recycling are well supported in studies of model fungi. The *S. cerevisiae* α-factor pheromone receptor, Ste2, is phosphorylated on the most distal serine/threonine residues on its C-terminal tail. Phosphomutation studies suggested that these residues are required for receptor-ligand sensitivity, revealing a regulatory role for this PTM. However, in subsequent truncation experiments, investigators demonstrated that removing the entire C-terminal tail of Ste2 resulted in a severe morphogenesis defect [57,58]. This observation is similar to our analysis of the phosphorylation site (T317A) of the Rra1 C-terminus. Mutating this residue inhibits the ability for Rra1 to localize in the plasma membrane, but does not inhibit its function, whereas removing the entire C-terminus renders the protein nonfunctional and the pathway inactive [16].

### Rim signaling regulates plasma membrane dynamics and Rra1 cycling

Although our experimental results support a model in which Rra1 localization in punctate structures at the cell surface is not necessary for activation, they also indicate an important link between the plasma membrane and *C. neoformans* Rim signaling. The question remains of why the *C. neoformans* Rra1 pH sensor localizes to the plasma membrane at low pH. Our previous work demonstrated that Rra1 functions similarly to the pH-sensing proteins in *S. cerevisiae* and *A. nidulans*. Specifically, these sensors use their C-terminal tails to sense changes in plasma membrane asymmetry and phospholipid distribution in order to efficiently responds to changes in extracellular pH [16,18,59–61]. Therefore, we suggest that the membrane localization of Rra1 allows for its condition-dependent internalization through the dynamics between its cytoplasmic tail and the phospholipids in the membrane. However, it is the internalized localization of this protein that allows it to interact with its downstream effectors and activate the Rim alkaline response.

The results from the work presented here further support the connection between Rim signaling and membrane dynamics through detailed lipidomics of the wildtype and *rim101*Δ mutant strains. This connection is further supported through rescue studies showing suppression of the *rim101*Δ pH-sensitive mutant phenotype when the growth media is supplemented with glycerol, the backbone of phospholipids. The inner leaflet of the fungal plasma membrane is enriched for specific bulky phospholipids, like phosphatidylserine (PS), whereas endosomes and vacuoles are not [55]. Our results showing increases in the PS levels of the *rim101*Δ mutant strain compared to wildtype at high pH, could represent altered integrity of both plasma and endosomal membranes and a disruption in the balance needed for proper protein cycling. The increased levels of PC also found in the *rim101*Δ mutant at high pH, also affect the ratio of membrane phospholipids, which might then affect protein trafficking throughout the cell. This altered membrane composition coupled with the results showing decreased ability for Rra1 to recycle back to the plasma membrane in the *rim101*Δ mutant, the T317A phosphomutant, and in conditions that affect overall membrane integrity support the dependence of Rra1 cycling on Rim-regulated membrane maintenance.

Additionally, it is known that Rim pathway outputs are involved in cell wall remodeling [3,5,9,16,62] and that specific membrane domain characteristics are dependent not only on lipid distribution and composition, but also on the proximity to the fungal cell wall [2,63]. Rim101 regulation of the fungal cell wall at high pH could have direct effects on cell wall turgor pressure and therefore would affect the shape and curvature of the plasma membrane allowing for membrane-associated proteins to establish themselves in microdomains. Furthermore, the significant increases in unsaturated and cumbersome PC and PS lipid species in the *rim101*Δ mutant strain might affect the formation of protein-localizing lipid rafts in the plasma membrane. These same species were significantly decreased in the wildtype strain in response to an increase in pH, further connecting the regulation of lipid ratios in the membrane, specifically proportions of unsaturated species, with the alkaline pH response.

The specific membrane microdomain localization of the *S. cerevisiae* pH sensors has been partially identified, and this identification might reveal insights regarding Rra1 membrane association. Previous investigators observed Rim21 localization as distinct from Membrane Compartment containing arginine permease Can1 (MCC) regions in the plasma membrane [64]. This is an important discovery because MCC domains cannot also function as sites of endocytosis due to their bulky nature and the inability for endocytosis machinery to assemble around cargo [65]. It has also been determined that Rim21 localizes to portions of the membrane that are devoid of cortical ER, eliminating MCL microdomains (Sterol transporter regions) as potential resting sites [66]. This is also an important distinction based on our previous studies that identified the sterol-mediated alkaline response as a Rim pathway-independent alkaline response process in *C. neoformans* [26].

We therefore conclude that these data support a model of alkaline pH-induced Rra1 internalization and recycling that intimately involve Rim-dependent membrane modifications as graphically depicted in Fig 7. In response to an alkaline shift, the *C. neoformans* Rra1 pH sensor is endocytosed through invagination of the plasma membrane where it resides in specific microdomains (1). The Nap1 adaptor protein stabilizes the Rra1 protein during this invagination through interaction with its cytosolic C-terminal tail [17] (2). The Rra1 protein, including its C-terminus, undergoes a conformational change to enable internalization and movement away from the plasma membrane allowing Rra1 to interact with downstream effectors. Once endocytosed, a clathrin coat forms around the Rra1-containing vesicle and the ESCRT machinery is recruited (3). Upstream Rim pathway components and downstream effectors (Rim23, Rim20, and the Rim13 protease) are then recruited to the plasma membrane as previously described [8] (4). This movement initiates cleavage of the terminal component of the Rim pathway, the Rim101 transcription factor (5a). Following cleavage, Rim101 translocates to the nucleus to aid in the transcription of virulence genes needed for growth of this fungus at alkaline pH, including genes involved in cell wall remodeling and membrane maintenance (5B). The clathrin-coated vesicle containing Rra1 is then coated with COPI and transported through the Golgi (6 & 7). This vesicle then sheds the COPI and clathrin coats and travels to the endoplasmic reticulum (ER) (8). When a decrease in pH is sensed, the Rra1 protein is then escorted from the ER (9) back through the Golgi where it is actively recoated with COPII coatomer and clathrin (10). The vesicle containing Rra1 is then transported back up to the plasma membrane to regions rich in sphingolipids and sterols (i.e. lipid rafts) (12). Rra1 then remains poised in the plasma membrane awaiting a shift in extracellular pH. Overall, these data help us to understand the role of the Rra1 pH-sensing protein in the Rim-dependent alkaline pH response and the mechanism by which it responds to extracellular stress in a relevant human fungal pathogen.

## Materials and Methods

### Strains, media, and growth conditions

Strains generated and used in these studies are shown in Table 3. Each phosphomutant and fluorescently tagged strain was generated in either the *C. neoformans* H99 *MATα* or the KN99 *MATa* genetic background. The MATa strain expressing Rra1-GFP (KMP81) was generated by a mating cross between the *MATα* strain expressing Rra1-GFP (KS310) and the *MATa* wildtype strain (KN99) (Table 3). Spores were selected on YPD medium + NAT/NEO and the ability to mate with *MATα* (*H99*). The *sre1*Δ + Rra1-GFP *MATa* (HEB99) and *rim101*Δ + Rra1-GFP *MATa* (HEB101) strains, were generated from a mating cross between KMP81 and the *sre1*Δ::NEO *MATα* (HEB5) and *rim101*Δ::NAT *MATα* (TOC35) strains, respectively (Table 3). Spores were selected for on YPD medium + NAT/NEO, the ability to mate with *MATα* (*H99*), and pH-sensitivity.

To generate all phosphomutant strains containing the GFP-tagged Rra1, *pKS85* (*pHIS3-RRA1-GFP-NAT*) plasmid was subjected to site directed mutagenesis to generate mutant alleles for each predicted phosphosite (described in more detail below). These mutated plasmids, listed in Table 4, were then biolistically transformed into the *rra1*Δ::*NEO* (KS336) full knockout strain (Table 3).

**Table 3.**
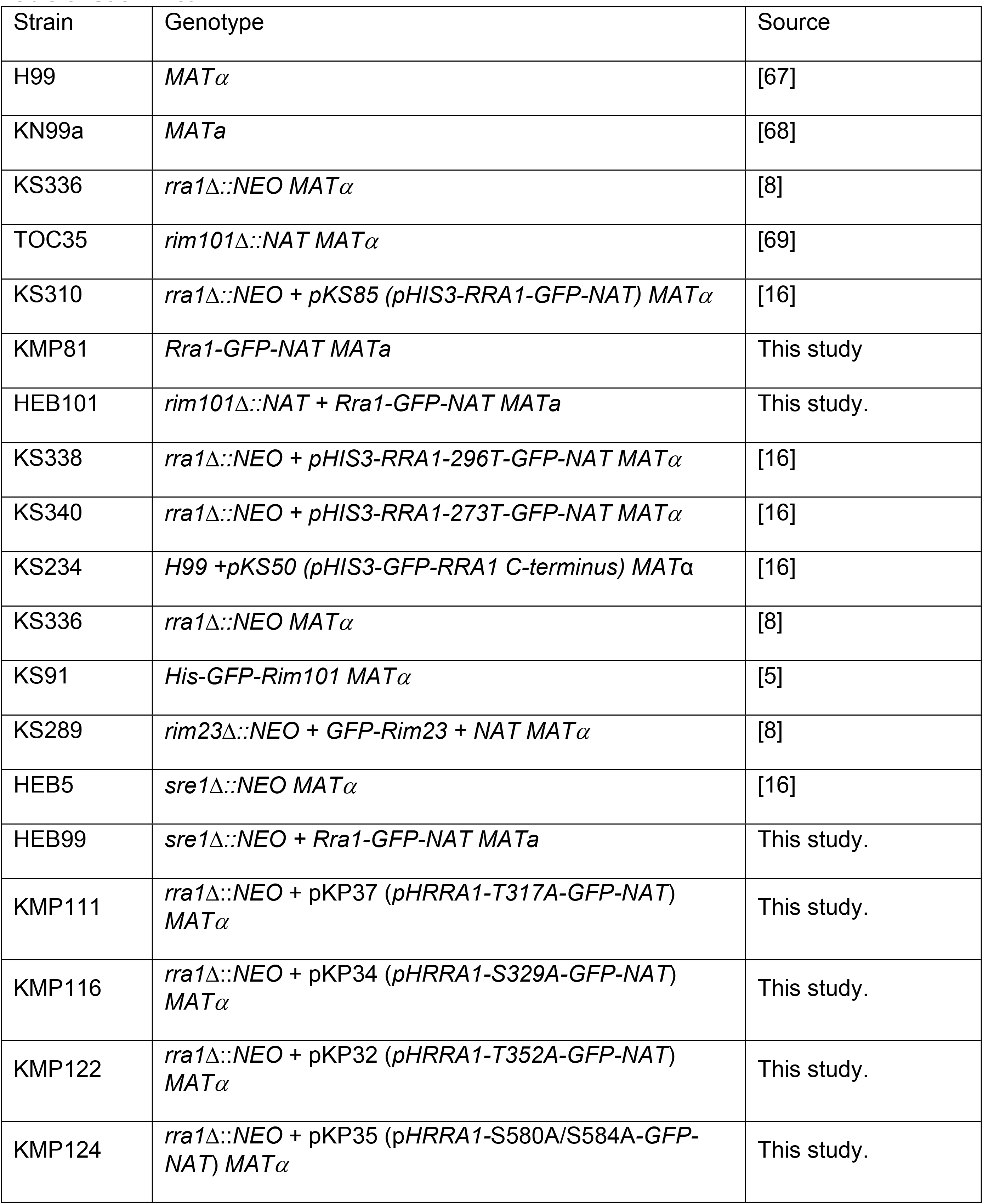
Strain List

**Table 4.** Plasmid List

| Plasmid | Open Reading Frame | Backbone | Reference |
| --- | --- | --- | --- |
| *pKP37* | *pHRR1-GFP-NAT T317A* | pHNAT | This study |
| *pKP34* | *pHRR1-GFP-NAT T329A* | pHNAT | This study |
| *pKP32* | *pHRR1-GFP-NAT T352A* | pHNAT | This study |
| *pKP35* | *pHRR1-GFP-NAT S580A/S584A* | pHNAT | This study |
| *pKS85* | *pHIS3-RRA1-GFP-NAT* | pHNAT | This study |

Strains were incubated in either Yeast Peptone Dextrose media (YPD) (1% yeast extract, 2% peptone, and 2% dextrose) or buffered media: YPD pH 4 and pH 8 media. Buffered media was made by adding 150 mM HEPES buffer to YPD, adjusting the pH with concentrated HCl (for pH 4) or NaOH (for pH 8.15), prior to autoclaving. 20% glucose was added to the media following sterilization and autoclaving. For YPD + glycerol plates, 0.4% glycerol was also added to the media following sterilization and autoclaving. For spot plate assays, strains were incubated overnight at 30°C with 150 rpm shaking in YPD, washed twice, resuspended in 1X PBS, and serially diluted onto selective media. Plates were then incubated at 30°C for 1-3 days and imaged.

### Microscopy

To analyze Rra1-GFP localization in various backgrounds, strains were incubated at 30°C for 18h with 150 rpm shaking in YPD. Cells were then pelleted and resuspended in either pH 4 or pH 8 Synthetic Complete media buffered with McIlvaine’s buffer [8]. For the FM4-64 (5 μg/μl; Invitrogen) colocalization studies, strains were grown overnight and shaken at 150 rpm and 30°C in YPD. Cells were pelleted, washed with PBS, and resuspended in 1 mL McIlvaine’s buffer pH 8. 1 μl of FM4-64 stock solution was added to cell suspension and cells were incubated on ice for 10 minutes and 20 minutes and imaged. Fluorescent images were captured using a Zeiss Axio Imager A1 fluorescence microscope equipped with an Axio-Cam MRM digital camera. Images were created using ImageJ software (Fiji) [70].

For Rra1 cycling microscopy, strains were incubated at 30°C with for 18h with 150 rpm shaking in YPD. Cells were then pelleted and resuspended in either pH 4 or pH 8 Synthetic Complete media buffered with McIlvaine’s buffer. Cells were then incubated for 60 minutes shaking at 30°C with shaking at 150 rpm. These cells were then pelleted, lightly resuspended, and imaged. Fluorescent images were captured as before. The cells that were grown in pH 8 McIlvaine’s buffer were re-pelleted and resuspended in pH 4 buffer and incubated for 30 minutes shaking at 30°C with 150 rpm. These cells were then pelleted, lightly resuspended, and imaged and are represented by the pH 8 to pH 4 images. Rra1-GFP localization studies in both the *sre1*Δ and T317A phosphomutant backgrounds was also performed using the same incubations (60 minutes in initial pH condition). For Rra1 cycling in the T317A phosphomutant background (Fig S3A), the same experiment was done but with shorter pre-incubations (30 minutes) in each extreme in order to see subtle phenotypes. Quantification of puncta per cell (2+) was done using ImageJ Software (Fiji) software [70] and a blinded identification of cells with membrane associated puncta in each condition as previously described [16]. Approximately 600 cells per condition/strain were analyzed. For Rra1 localization and cycling in the *rim101*Δ and *sre1*Δ mutant backgrounds, the Rra1-GFP *MATa* strain (KMP81) was used as the positive control. For Rra1 localization and cycling in the phosphomutant backgrounds, the Rra1-GFP *MATα* strain (KS310) was used.

For Rra1-GFP (KS310) and GFP-Rim101 (KS91) localization with Pitstop-2 (Sigma) treatment experiments, strains were incubated in YPD at 30°C for 18h with 150 rpm shaking. Cells were then pelleted and resuspended in either pH 4 or pH 8 McIlvaine’s buffer for 10 minutes following treatment with 20 μM Pitstop-2 (Rra1-GFP experiment) or both 20 and 42 μM Pitstop-2 (eGFP-Rim101 experiment) or vehicle control (DMSO). Cells were treated and incubated at 37°C with shaking at 150 rpm. For Rra1 localization, the mean values and standard errors of cells with clear endomembrane localization at pH 8 was quantified using ImageJ software (Fiji) (~600 cells/condition; 4 biological replicates). Quantification graphs and statistics using a student’s t-test were generated in GraphPad Prism (GraphPad Prism version 8.00 for Mac, GraphPad Software, San Diego California USA, www.graphpad.com). For Rim101 localization, the mean values and standard errors of cells with clear nuclear localization at either pH in increasing amounts of drug were quantified using Fiji (~600 cells/condition; 3 biological replicates). GraphPad Prism software was used to generate the graph and the One-way ANOVA, Tukey’s multiple comparison statistical analyses.

### Drug Susceptibility Tests

Pitstop-2 treatment experiments to determine susceptibility of wildtype *C. neoformans* (H99) cells to this treatment at varying pH was performed by broth microdilution. Specifically, cells were incubated at 30°C for 18h with 150 rpm shaking in YPD. Pistop-2 resuspended in DMSO was serially diluted in Synthetic Complete media buffered to pH 6.6, 6.8, 7, 7.2, or 7.4 with McIlvaine’s buffer. Fungal cells were then normalized and diluted in Synthetic Complete media buffered to the same pH values and added to the corresponding pH well containing Pitstop-2. Plates were incubated at 30°C for 72 hours, and the MIC was determined to be the lowest concentration of drug that led to no fungal cell growth.

Filipin III treatment experiments were carried out similarly. Wildtype cells (H99) were treated with increasing concentrations of Filipin III, which had been serially diluted in Synthetic Complete media buffered to pH 4, 5, 6, 7, and 8. Fungal cells were, again, normalized and diluted in the same media and added to wells containing Filipin III. Plates were incubated at 30°C for 48 hours, and the MIC was determined to be the lowest concentration of drug that led to no fungal cell growth.

### Protein Extraction, Immunoprecipitation, and Western Blot

Protein extracts for the protein interaction studies were prepared as in a similar manner to that previously described [8,16,17]. Briefly, the wildtype (H99 untagged strain), the GFP-Rim23 (KS289), and the GFP-Rra1-Ct (KS234) strains were incubated at 30 °C for 18h with 150 rpm shaking in YPD pH 4. Cells were then pelleted and resuspended in either pH 4 again or switched to YPD pH 8. These cells were incubated for 1 hour and immediately pelleted and flash frozen. Cells were then lysed using 0.4 mL lysis buffer containing 2x protease inhibitors (Complete, Mini, EDTA-free; Roche), 1x phosphatase inhibitors (PhosStop; Roche) and 1 mM phenylmethanesulfonyl-fluoride (PMSF). Lysis was performed by bead beating (0.5 mL of 3 μM glass beads in a Mini-BeadBeater-16 (BioSpec), 6 cycles of 30 seconds each with a one-minute ice incubation between bead-beating cycle). Supernatants were transferred to new tubes and washed 3 times with 0.4 mL of lysis buffer. The crude pellet was then pelleted through centrifugation at 15,000 rpm, 4 °C, for 5 minutes, and the supernatant (cell lysate) was transferred to a new tube and further ultracentrifuged at 100,000 x g. Proteins were immunoprecipitated by the addition of 50 μl pre-equilibrated GFP-Trap resin (Chromotek) and inverted for 2 hours at 4 °C. Mass spectrometry experiments were performed at an *n* of 1 by the Duke Proteomics Core Facility as previously described [17].

For Rra1-GFP protein gels and western blot analysis, the same experimental procedure as above was performed, but with some modifications. Briefly, the Rra1-GFP (KS310), the Rra1-GFP 296T truncation mutant (KS336) and the Rra1-GFP T317A phosphomutant (KMP111) were grown at 30 °C for 18h with 150 rpm shaking in YPD pH 4. Cells were then pelleted and resuspended in YPD pH 8 and incubated for 1.5 hours prior to lysis and protein extraction as outlined above. These lysates were not subjected to GFP-Trap pull down, instead whole cell lysate protein concentrations were measured using bicinchoninic acid assay (BCA) and protein samples were normalized and diluted in 4X NuPage lithium dodecyl sulfate (LDS) loading buffer and 10X NuPage Reducing Agent to a 1X concentration and boiled at 100°C for 5 mins. Western blots were performed as described previously using a 4-12% NuPage BisTris gel. To probe and detect Rra1-GFP, immunoblots were incubated in anti-GFP primary antibody (using a 1/10,000 dilution, Roche) and then in secondary anti-mouse peroxidase-conjugated secondary antibody (using a 1/25,000 dilution, Jackson Labs). Proteins were detected by enhanced chemiluminescence (ECL Prime Western blotting detection reagent; GE Healthcare).

### GO term analysis (FungiFun, FunCat, and Cellular Compartment GO)

The interactomes of the GFP-Rra1-Ct (KS234) and GFP-Rim23 (KS289) (extracted and analyzed as above) were run through FungiFun software to determine significantly enriched Gene Ontology (GO) categories [24]. The interactomes of these proteins at pH 8 were compared to that of the non-tagged wildtype control in the same condition and interactors with at least 5 exclusive peptides that were only present in the GFP-tagged sample were prioritized. This prioritization excluded potential false-positive interactions. Through the FungiFun program, the interactomes were analyzed by Funcat to observe general categories and cellular processes that are enriched in these data sets using the following parameters: hypergeometric distribution, p-value of 0.05, overrepresentation (enrichment), Benjamini-Hochberg procedure, and directly annotated associations. These data were also run through GO-term analysis looking specifically at cellular compartments to observe specific cellular locations that are significantly represented in these interactomes. The cellular component analysis was run using the same parameters. The specific CNAG #s and gene names in each category are in Tables 1 and 2, and the full interactomes are in Table S1.

### Phosphoproteomics

Protein for the phosphoproteomics experiment was harvested from cells prepared as described above for the protein interaction studies. Cells from the KS310 strain expressing Rra1-GFP were lysed also as described above, but with the addition of 1X PhosStop phosphatase inhibitor (Roche). After lysis, crude lysates were cleared at 5000 rpm for 10 minutes. Protein was then normalized such that immunoprecipitation was performed on 5 mg of protein per sample. Immunoprecipitation was performed as described above. Samples were submitted to the Duke University Proteomics core. For Rra1-GFP samples, samples were divided and part treated as described above for mass spectrometry, and part subjected to TiO2 enrichment of phosphopeptides after digestion and before mass spectrometry.

### Site-directed mutagenesis/Phosphomutant generation

To create the non-phosphorylated site mutants, the *pHRRA1-GFP* (pKS85) (Table 4) plasmid was PCR-amplified with Phusion HF Polymerase (NEB), using primers designed using the QuikChange Primer Design tool (Agilent). Site-directed mutagenesis primers can be found in Table 5. PCR products were PCR purified using the DNA Clean and Concentrator kit (Zymo Research), then transformed into One Shot TOP10 competent cells (ThermoFisher Scientific). Each mutant construct was sequenced to ensure that no unintended mutations were introduced, and subsequently transformed into the *rra1*Δ mutant strain (KS336). Using quantitative real time-PCR, we identified and prioritized transformants in which each allele was expressed at levels similar to the *RRA1-GFP* control *RRA1* primers listed in Table 5.

**Table 5.**
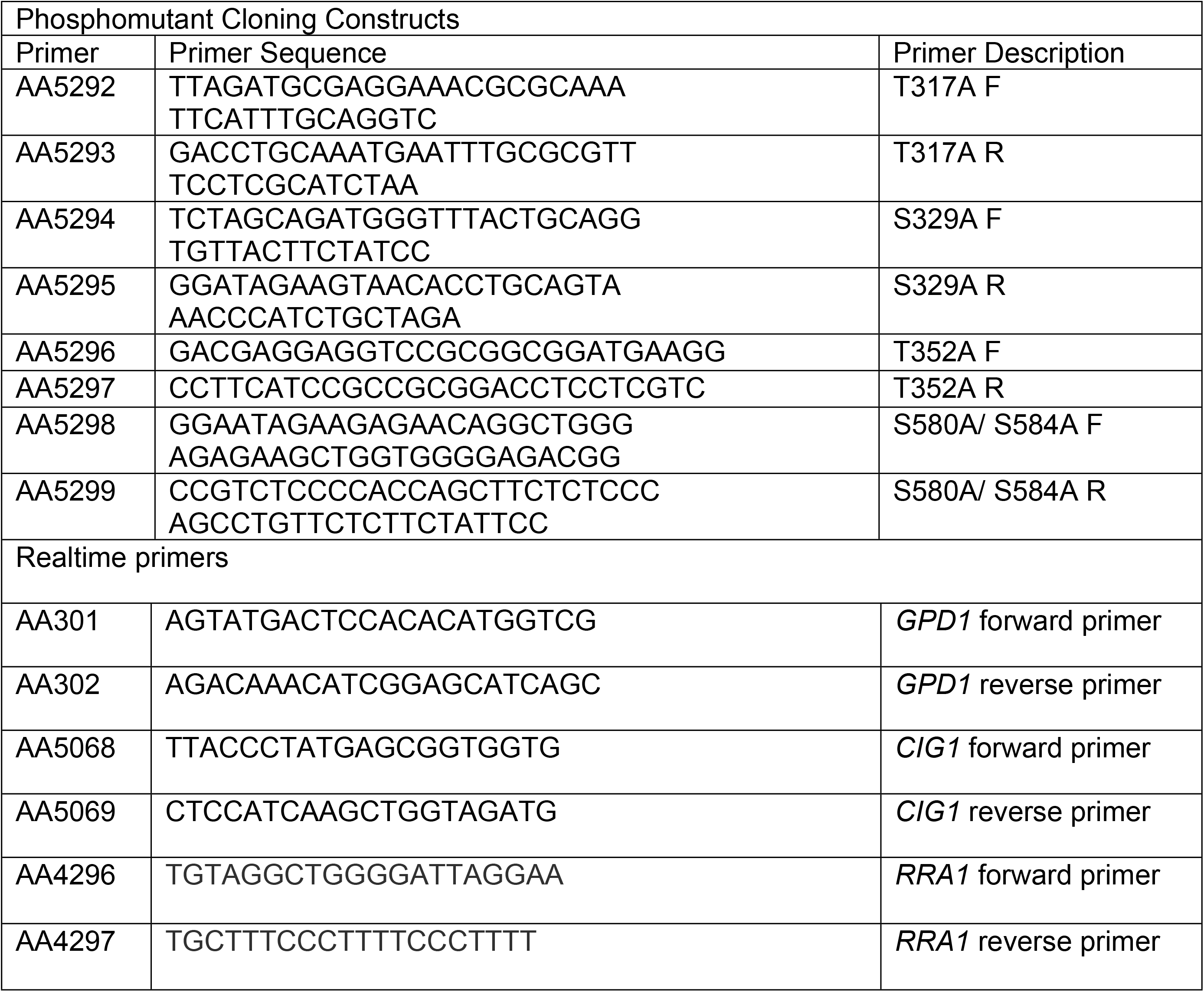
Primer List

### RNA Extraction and Quantitative Real Time PCR

qRT-PCR was performed on the T317A phosphomutant as previously described [16]. Briefly, three biological replicates of wildtype (H99), *rra1*Δ (KS336)*, rra1*Δ +Rra1-GFP (KS310), and *rra1*Δ + Rra1-GFP T317A (KMP111) were prepped and RNA-extracted. Strains were incubated overnight at 30 °C for 18h with 150 rpm shaking in YPD media. Cells were pelleted and resuspended in YPD pH 8 media and incubated for 1.5 hours at 30 °C with 150 rpm shaking. Cells were then pelleted, flash frozen on dry ice, and lyophilized overnight. RNA was extracted by using the Qiagen RNeasy Plant Minikit with on column DNase digestion (Qiagen, Valencia, CA). cDNA was prepped by reverse-transcriptase PCR using the AffinityScript cDNA QPCR Synthesis kit (Agilent Technologies. qRT-PCR reactions were performed as previously described [16,71] using *RRA1* and *CIG1* primers listed in Table 5.

### Lipidomics Analysis

The lipid extraction was performed as described in [72]. Briefly, *C. neoformans* strains were grown in minimal media (100 mM HEPES, 0.67 % YNB without amino acids, 2 % glucose, pH 4 or 8) at 30 °C for 48h under agitation. Cell suspensions were centrifuged at 1,734 g for 10 minutes, the supernatant was removed, and pellet was washed twice with milliQ water. Cultures were counted in a hemocytometer and 5×10^8^ cells per sample were transferred to a glass tube. The suspensions were centrifuged again, and supernatant was removed. Then, each sample was suspended in 1.5 ml of Mandala buffer and vortexed vigorously for 20 seconds. The extraction was performed as described in [73], followed by Bligh and Dyer extraction [74]. A quarter of each sample obtained from the Bligh and Dyer Extraction was reserved for inorganic phosphate (Pi) determination, so the relative phospholipid signal was normalized by the Pi abundance. The organic phase was transferred to a new tube, dried and used for MS analysis.

## Acknowledgements

We acknowledge the Duke Proteomics Shared Resource for their assistance with the various projects in this study. We acknowledge and thank Sandra Breeding for her assistance with media and reagents. Finally, we acknowledge our NIH funding (R01 AI074677, 1F31A140427-01A1, P01 AI104533, to JAA and R01225770 to MDP) that made these studies possible. Dr. Del Poeta is a co-founder and chief scientific officer of MicroRid Technologies Inc. All authors have no relevant conflicts of interest.

## Author Contributions

HEB, CMF, KMP, MDP, and JAA were involved with the conception and design of experiments and the writing process. HEB, KMP, CMF, and KM were involved in the acquisition of the data. All authors participated in the analysis and interpretation of the data.

---

**Fig S1. Rra1 membrane puncta are unaffected by inhibition of clathrin-mediated endocytosis at low pH**

A. pH-dependent localization of the Rra1-GFP fusion protein in response to pH 4 and pH 8 McIlvaine’s media for 10 minutes following treatment with either 20 μM Pitstop-2 or DMSO. GFP signal was assessed by epifluorescence microscopy (Zeiss Axio Imager A1) using the appropriate filter. Long white arrows indicate cells with > 2 membrane puncta (MP), Asterisks indicate cells containing intermediate localization, and short white arrows indicate clear endomembrane/ER localization. White scale bars indicate 5 microns.

B. Quantification of Rra1-GFP localization at pH 4 and pH 8. The mean values and standard errors of cells with clear ER localization at pH 8 was quantified using ImageJ software (Fiji) (~600 cells/condition; 4 biological replicates). One-way ANOVA Tukey’s multiple comparison test: No significant difference (ns) between Rra1 MP at pH 4 between untreated and treated groups.

**Fig S2. Predictive modeling of the Rra1 protein**

Prediction software Itasser (Iterative Threading ASSEmbly Refinement) used a hierarchical approach to model the most likely configurations of the Rra1 protein according to its amino acid sequence. (A) represents the first predicted model of Rra1 and has a C-score of −1.70 (range between −5 and 2, with higher scores representing higher confidence). (B) represents the second most likely modeled configuration of the protein, with a lower C-score of −2.31. Both secondary structures with C-terminal tails highlighted in teal and space fill models are shown for each predicted model (generated in Protean 3D software using Itasser predictions).

C. Schematic of the Rra1 protein functional domains and PONDR predicted disordered regions. Rainbow colors correspond with the 7 predicted transmembrane domains in the space-filled models in (A) and (B) as well as with the transmembrane domain prediction as analyzed by Protean 3D software using Von Heijne modeling in (D). A separate disorder prediction software (JRONN) was also run through Protean 3D revealing the highly disordered C-terminal tail (E).

**Fig S3. Rra1 cycling and pathway activation in the T317A mutant**

A. pH-dependent localization and recycling of the Rra1 protein GFP fusion construct in response to pH 4 and pH 8 McIlvaine’s media for 30 minutes and then back to pH 4 media for 30 minutes in the wildtype and T317A phosphomutant backgrounds. GFP signal was assessed by epifluorescence microscopy (Zeiss Axio Imager A1) using the appropriate filter. White scale bars indicate 5 microns. Quantification of Rra1-GFP localization at pH 4 and pH 8. The mean values and standard errors of cells with > 2 membrane puncta formed at pH 4 and 8 was quantified using ImageJ software (Fiji) (~600 cells/condition; 3 biological replicates). One-way ANOVA Tukey’s multiple comparison test: *** = p = 0.0009

Quantitative Realtime PCR analysis of (B) *RRA1* and (C) a known Rim pathway output, *CIG1*, in the wildtype, *rra1Δ, rra1*Δ + Rra1-GFP, and *rra1*Δ + Rra1-GFP + T317A phosphomutant. Strains were incubated in YPD pH 8 conditions for 1.5 hours prior to RNA-extraction (in biological triplicate) and analysis of transcript abundance by PCR. log_2_ fold change of *CIG1* expression of the various strains is indicated compared to wildtype.

**Table S1:** Proteomics data.

Page 1. List of the prioritized GFP-Rra1-Ct interactors at pH 8 compared to untagged control.

Page 2. List of prioritized GFP-Rra1-Ct interactors with assigned FunCat categories.

Page 3. List of prioritized GFP-Rra1-Ct interactors with assigned GO Cellular Compartment categories.

Page 4. List of the prioritized Rim23-GFP interactors at pH 8 compared to untagged control.

Page 5. List of prioritized Rim23-GFP interactors with assigned FunCat categories.

Page 6. List of prioritized Rim23-GFP interactors with assigned GO Cellular Compartment categories.

**Table S2: Phospholipidomics data**

Phosphatidylcholine (page 1), phosphatidylserine (page 2), and phosphatidylethanolamine (page

3) lipid species in the wildtype, *rim101*Δ, and *rim101*Δ + *RIM101* strains at pH 4 and pH 8.

## References

1. Brakhage AA, Spröte P, Al-Abdallah Q, Gehrke A, Plattner H, Tüncher A. Regulation of penicillin biosynthesis in filamentous fungi. Advances in biochemical engineering/biotechnology. Adv Biochem Eng Biotechnol; 2004. pp. 45–90. doi:10.1007/b99257

2. Athanasopoulos A, André B, Sophianopoulou V, Gournas C. Fungal plasma membrane domains. FEMS Microbiology Reviews. Oxford University Press; 2019. pp. 642–673. doi:10.1093/femsre/fuz022

3. Mira NP, LourenÃ§o AB, Fernandes AR, Becker JD, SÃi-Correia I. The RIM101 pathway has a role in *Saccharomyces cerevisiae* adaptive response and resistance to propionic acid and other weak acids. FEMS Yeast Res. 2009;9: 202–216. doi:10.1111/j.1567-1364.2008.00473.x

4. O’Meara TR, Holmer SM, Selvig K, Dietrich F, Alspaugh JA. *Cryptococcus neoformans* Rim101 is associated with cell wall remodeling and evasion of the host immune responses. MBio. 2013;4. doi:10.1128/mBio.00522-12

5. O’Meara TR, Xu W, Selvig KM, O’Meara MJ, Mitchell AP, Alspaugh JA. The *Cryptococcus neoformans* Rim101 transcription factor directly regulates genes required for adaptation to the host. Mol Cell Biol. 2014;34: 673–84. doi:10.1128/MCB.01359-13

6. Sherrington SL, Sorsby E, Mahtey N, Kumwenda P, Lenardon MD, Brown I, et al. Adaptation of *Candida albicans* to environmental pH induces cell wall remodelling and enhances innate immune recognition. Noverr MC, editor. PLOS Pathog. 2017;13: e1006403. doi:10.1371/journal.ppat.1006403

7. Cornet M, Gaillardin C. pH signaling in human fungal pathogens: a new target for antifungal strategies. Eukaryot Cell. 2014;13: 342–52. doi:10.1128/EC.00313-13

8. Ost KS, O’Meara TR, Huda N, Esher SK, Alspaugh JA. The *Cryptococcus neoformans* Alkaline Response Pathway: Identification of a Novel Rim Pathway Activator. PLoS Genet. 2015;11. doi:10.1371/journal.pgen.1005159

9. Ost KS, Esher SK, Leopold Wager CM, Walker L, Wagener J, Munro C, et al. Rim Pathway-Mediated Alterations in the Fungal Cell Wall Influence Immune Recognition and Inflammation. MBio. 2017;8: e02290–16. doi:10.1128/mBio.02290-16

10. Hommel B, Mukaremera L, Cordero RJB, Coelho C, Desjardins CA, Sturny-Leclère A, et al. Titan cells formation in *Cryptococcus neoformans* is finely tuned by environmental conditions and modulated by positive and negative genetic regulators. PLoS Pathog. 2018;14: e1006982. doi:10.1371/journal.ppat.1006982

11. Selvig K, Alspaugh JA. pH Response Pathways in Fungi: Adapting to Host-derived and Environmental Signals. Mycobiology. 2011;39: 249–56. doi:10.5941/MYCO.2011.39.4.249

12. Bertuzzi M, Schrettl M, Alcazar-Fuoli L, Cairns TC, Muñoz A, Walker LA, et al. The pH-responsive PacC transcription factor of *Aspergillus fumigatus* governs epithelial entry and tissue invasion during pulmonary aspergillosis. PLoS Pathog. 2014;10: e1004413. doi:10.1371/journal.ppat.1004413

13. Davis D, Edwards JE, Mitchell AP, Ibrahim AS. *Candida albicans* RIM101 pH response pathway is required for host-pathogen interactions. Infect Immun. 2000;68: 5953–9. Available: http://www.ncbi.nlm.nih.gov/pubmed/10992507

14. Antonio C-CJ, Lucila O-C, Miriam T-S, Scott G, José R-H. Functional analysis of the pH responsive pathway Pal/Rim in the phytopathogenic basidiomycete *Ustilago maydis*. Fungal Genet Biol. 2010;47. doi:10.1016/j.fgb.2010.02.004

15. Rajasingham R, Smith RM, Park BJ, Jarvis JN, Govender NP, Chiller TM, et al. Global burden of disease of HIV-associated cryptococcal meningitis: an updated analysis. Lancet Infect Dis. 2017;17: 873–881. doi:10.1016/S1473-3099(17)30243-8

16. Brown HE, Ost KS, Esher SK, Pianalto KM, Saelens JW, Guan Z, et al. Identifying a Novel Connection Between the Fungal Plasma Membrane and pH-Sensing. Mol Microbiol. 2018 [cited 10 Jul 2018]. doi:10.1111/mmi.13998

17. Pianalto KM, Ost KS, Brown HE, Alspaugh JA. Characterization of additional components of the environmental pH-sensing complex in the pathogenic fungus *Cryptococcus neoformans*. J Biol Chem. 2018;293: 9995–10008. doi:10.1074/jbc.RA118.002741

18. Nishino K, Obara K, Kihara A. The C-terminal Cytosolic Region of Rim21 Senses Alterations in Plasma Membrane Lipid Composition: Insights into sensing mechanisms for plasma membrane lipid asymmetry. J Biol Chem. 2015;290: 30797–805. doi:10.1074/jbc.M115.674382

19. Fernandes CM, Goldman GH, Del Poeta M. Biological Roles Played by Sphingolipids in Dimorphic and Filamentous Fungi. 2018. Available: http://mbio.asm.org/

20. Munshi MA, Gardin JM, Singh A, Luberto C, Rieger R, Bouklas T, et al. The Role of Ceramide Synthases in the Pathogenicity of *Cryptococcus neoformans*. Cell Rep. 2018;22: 1392–1400. doi:10.1016/j.celrep.2018.01.035

21. Luberto C, Toffaletti DL, Wills EA, Tucker SC, Casadevall A, Perfect JR, et al. Roles for inositol-phosphoryl ceramide synthase 1 (IPC1) in pathogenesis of *C. neoformans*. Genes Dev. 2001;15: 201–12. doi:10.1101/gad.856001

22. Lev S, Rupasinghe T, Desmarini D, Kaufman-Francis K, Sorrell TC, Roessner U, et al. The PHO signaling pathway directs lipid remodeling in *Cryptococcus neoformans* via DGTS synthase to recycle phosphate during phosphate deficiency. Bahn Y-S, editor. PLoS One. 2019;14: e0212651. doi:10.1371/journal.pone.0212651

23. Hu G, Caza M, Cadieux B, Chan V, Liu V, Kronstad J. *Cryptococcus neoformans* requires the ESCRT protein Vps23 for iron acquisition from heme, for capsule formation, and for virulence. Infect Immun. 2013;81: 292–302. doi:10.1128/IAI.01037-12

24. Priebe S, Kreisel C, Horn F, Guthke R, Linde J. FungiFun2: A comprehensive online resource for systematic analysis of gene lists from fungal species. Bioinformatics. 2015;31: 445–446. doi:10.1093/bioinformatics/btu627

25. Telzrow CL, Nichols CB, Castro-Lopez N, Wormley FL, Alspaugh JA. A fungal arrestin protein contributes to cell cycle progression and pathogenesis. MBio. 2019;10. doi:10.1128/mBio.02682-19

26. Brown HE, Telzrow CL, Saelens JW, Fernandes L, Alspaugh JA. Sterol-Response Pathways Mediate Alkaline Survival in Diverse Fungi. MBio. 2020;11. doi:10.1128/mBio.00719-20

27. Chung D, Barker BM, Carey CC, Merriman B, Werner ER, Lechner BE, et al. ChIP-seq and In Vivo Transcriptome Analyses of the *Aspergillus fumigatus* SREBP SrbA Reveals a New Regulator of the Fungal Hypoxia Response and Virulence. Doering TL, editor. PLoS Pathog. 2014;10: e1004487. doi:10.1371/journal.ppat.1004487

28. Bien CM, Chang YC, Nes WD, Kwon-Chung KJ, Espenshade PJ. *Cryptococcus neoformans* Site-2 protease is required for virulence and survival in the presence of azole drugs. Mol Microbiol. 2009;74: 672–690. doi:10.1111/j.1365-2958.2009.06895.x

29. Chang YC, Ingavale SS, Bien C, Espenshade P, Kwon-Chung KJ. Conservation of the sterol regulatory element-binding protein pathway and its pathobiological importance in *Cryptococcus neoformans*. Eukaryot Cell. 2009;8: 1770–9. doi:10.1128/EC.00207-09

30. Yang J, Zhang Y. I-TASSER server: New development for protein structure and function predictions. Nucleic Acids Res. 2015;43: W174–W181. doi:10.1093/nar/gkv342

31. Yang J, Yan R, Roy A, Xu D, Poisson J, Zhang Y. The I-TASSER suite: Protein structure and function prediction. Nature Methods. Nature Publishing Group; 2014. pp. 7–8. doi:10.1038/nmeth.3213

32. Roy A, Kucukural A, Zhang Y. I-TASSER: A unified platform for automated protein structure and function prediction. Nat Protoc. 2010;5: 725–738. doi:10.1038/nprot.2010.5

33. Garner E RPDABC and OZ. Predicting binding regions within disordered proteins. Genome Informatics. 1999;10: 41–50. Available: http://www.pondr.com/pondr-tut5.html

34. Dunker AK, Lawson JD, Brown CJ, Williams RM, Romero P, Oh JS, et al. Intrinsically disordered protein. J Mol Graph Model. 2001;19: 26–59. doi:10.1016/S1093-3263(00)00138-8

35. Epp E, Nazarova E, Regan H, Douglas LM, Konopka JB, Vogel J, et al. Clathrin-and arp2/3-independent endocytosis in the fungal pathogen *Candida albicans*. MBio. 2013;4. doi:10.1128/mBio.00476-13

36. De Duve C. The origin of eukaryotes: A reappraisal. Nature Reviews Genetics. Nature Publishing Group; 2007. pp. 395–403. doi:10.1038/nrg2071

37. Conner SD, Schmid SL. Regulated portals of entry into the cell. Nature. Nature Publishing Group; 2003. pp. 37–44. doi:10.1038/nature01451

38. Miaczynska M, Stenmark H. Mechanisms and functions of endocytosis. Journal of Cell Biology. The Rockefeller University Press; 2008. pp. 7–11. doi:10.1083/jcb.200711073

39. Gonzá Lez-Gaitá N M, Stenmark H. Endocytosis and Signaling: A Relationship under Development. Cell. 2003.

40. Park Y-D, Chen SH, Camacho E, Casadevall A, Williamson PR. Role of the ESCRT Pathway in Laccase Trafficking and Virulence of *Cryptococcus neoformans*. 2020 [cited 6 Aug 2020]. doi:10.1128/IAI.00954-19

41. Hurley JH, Emr SD. The ESCRT complexes: Structure and mechanism of a membrane-trafficking network. Annual Review of Biophysics and Biomolecular Structure. NIH Public Access; 2006. pp. 277–298. doi:10.1146/annurev.biophys.35.040405.102126

42. Weissman Z, Shemer R, Conibear E, Kornitzer D. An endocytic mechanism for haemoglobin-iron acquisition in *Candida albicans*. Mol Microbiol. 2008;69: 201–217. doi:10.1111/j.1365-2958.2008.06277.x

43. Park Y-D, Hui Chen S, Camacho E, Casadevall A. Role of the ESCRT pathway in Laccase Trafficking and Virulence of *Cryptococcus neoformans*. 2020 [cited 21 Apr 2020]. doi:10.1128/IAI.00954-19

44. Kamzolkina V V, Kiselica MA, Kudryavtseva OA, Shtaer O V, Mazheika IS. Endocytosis and Its Inhibitors in Basidiomycetous Fungus *Rhizoctonia solani*. Seriya. 2017;72: 149–157. doi:10.3103/S0096392517030063

45. Wang P, Shen G. The endocytic adaptor proteins of pathogenic fungi: Charting new and familiar pathways. Medical Mycology. NIH Public Access; 2011. pp. 449–457. doi:10.3109/13693786.2011.553246

46. Nikko E, Pelham HRB. Arrestin-mediated endocytosis of yeast plasma membrane transporters. Traffic. 2009;10: 1856–1867. doi:10.1111/j.1600-0854.2009.00990.x

47. Barwell KJ, Boysen JH, Xu W, Mitchell AP. Relationship of DFG16 to the Rim101p pH response pathway in *Saccharomyces cerevisiae* and *Candida albicans*. Eukaryot Cell. 2005;4: 890–899. doi:10.1128/EC.4.5.890-899.2005

48. Maeda T. The signaling mechanism of ambient pH sensing and adaptation in yeast and fungi. FEBS J. 2012;279: 1407–1413. doi:10.1111/j.1742-4658.2012.08548.x

49. Xu W, Smith FJ, Subaran R, Mitchell AP. Multivesicular body-ESCRT components function in pH response regulation in *Saccharomyces cerevisiae* and *Candida albicans*. Mol Biol Cell. 2004;15: 5528–5537. doi:10.1091/mbc.E04-08-0666

50. Boysen JH, Mitchell AP. Control of Bro1-domain protein Rim20 localization by external pH, ESCRT machinery, and the *Saccharomyces cerevisiae* Rim101 pathway. Mol Biol Cell. 2006;17: 1344–1353. doi:10.1091/mbc.E05-10-0949

51. Lucena-Agell D, Galindo A, Arst HN, Peñalva MA. *Aspergillus nidulans* ambient pH signaling does not require endocytosis. Eukaryot Cell. 2015;14: 545–553. doi:10.1128/EC.00031-15

52. Galindo A, Hervás-Aguilar A, Rodríguez-Galán O, Vincent O, Arst HN, Tilburn J, et al. PalC, one of two Bro1 domain proteins in the fungal pH signalling pathway, localizes to cortical structures and binds Vps32. Traffic. 2007;8: 1346–1364. doi:10.1111/j.1600-0854.2007.00620.x

53. Galindo A, Calcagno-Pizarelli AM, Arst HN, Peñalva MÁ. An ordered pathway for the assembly of fungal ESCRT-containing ambient pH signalling complexes at the plasma membrane. J Cell Sci. 2012;125: 1784–1795. doi:10.1242/jcs.098897

54. Dimou S, Diallinas G. Life and Death of Fungal Transporters under the Challenge of Polarity. International journal of molecular sciences. NLM (Medline); 2020. p. 5376. doi:10.3390/ijms21155376

55. Ma M, Burd CG. Retrograde trafficking and plasma membrane recycling pathways of the budding yeast *Saccharomyces cerevisiae*. Traffic. 2020;21: 45–59. doi:10.1111/tra.12693

56. Woods B, Lai H, Wu CF, Zyla TR, Savage NS, Lew DJ. Parallel Actin-Independent Recycling Pathways Polarize Cdc42 in Budding Yeast. Curr Biol. 2016;26: 2114–2126. doi:10.1016/j.cub.2016.06.047

57. Davis C, Dube P, Konopka JB. Afr1p Regulates the *Saccharomyces cerevisiae*-Factor Receptor by a Mechanism That Is Distinct From Receptor Phosphorylation and Endocytosis. Genetics. 1998.

58. Chen Q, Konopka JB. Regulation of the G-Protein-Coupled-Factor Pheromone Receptor by Phosphorylation. Mol Cell Biol. 1996. Available: http://mcb.asm.org/

59. Peñalva MA, Lucena-Agell D, Arst HN. Liaison alcaline: Pals entice non-endosomal ESCRTs to the plasma membrane for pH signaling. Curr Opin Microbiol. 2014;22: 49–59. doi:10.1016/J.MIB.2014.09.005

60. Obara K, Kihara A. Signaling events of the Rim101 pathway occur at the plasma membrane in a ubiquitination-dependent manner. Mol Cell Biol. 2014;34: 3525–34. doi:10.1128/MCB.00408-14

61. Lucena-Agell D, Hervás-Aguilar A, Múnera-Huertas T, Pougovkina O, Rudnicka J, Galindo A, et al. Mutational analysis of the *Aspergillus* ambient pH receptor PalH underscores its potential as a target for antifungal compounds. Mol Microbiol. 2016;101: 982–1002. doi:10.1111/mmi.13438

62. Brown HE, Esher SK, Alspaugh JA. Chitin: A “Hidden Figure” in the Fungal Cell Wall. 2019. doi:10.1007/82_2019_184

63. Farnoud AM, Bryan AM, Kechichian T, Luberto C, Poeta M Del. The Granuloma Response Controlling Cryptococcosis in Mice Depends on the Sphingosine Kinase 1-Sphingosine 1-Phosphate Pathway. 2015 [cited 25 Feb 2020]. doi:10.1128/IAI.00056-15

64. Obara K, Kihara A. The Rim101 pathway contributes to ER stress adaptation through sensing the state of plasma membrane. Biochem J. 2017;474: 51–63. doi:10.1042/BCJ20160580

65. Brach T, Godlee C, Moeller-Hansen I, Boeke D, Kaksonen M. The initiation of clathrin-mediated endocytosis is mechanistically highly flexible. Curr Biol. 2014;24: 548–554. doi:10.1016/j.cub.2014.01.048

66. Obara K, Yamamoto H, Kihara A. Membrane protein Rim21 plays a central role in sensing ambient pH in *Saccharomyces cerevisiae*. J Biol Chem. 2012;287: 38473–38481. doi:10.1074/jbc.M112.394205

67. Perfect JR, Lang SD, Durack DT. Chronic cryptococcal meningitis: a new experimental model in rabbits. Am J Pathol. 1980;101: 177–94. Available: http://www.ncbi.nlm.nih.gov/pubmed/7004196

68. Nielsen K, Cox GM, Wang P, Toffaletti DL, Perfect JR, Heitman J. Sexual cycle of *Cryptococcus neoformans* var. grubii and Virulence of congenic a and α isolates. Infect Immun. 2003;71: 4831–4841. doi:10.1128/IAI.71.9.4831-4841.2003

69. O’Meara TR, Norton D, Price MS, Hay C, Clements MF, Nichols CB, et al. Interaction of *Cryptococcus neoformans* Rim101 and protein kinase A regulates capsule. PLoS Pathog. 2010;6: e1000776. doi:10.1371/journal.ppat.1000776

70. Schindelin J, Arganda-Carreras I, Frise E, Kaynig V, Longair M, Pietzsch T, et al. Fiji: an open-source platform for biological-image analysis. Nat Methods. 2012;9: 676–682. doi:10.1038/nmeth.2019

71. Cramer KL, Gerrald QD, Nichols CB, Price MS, Alspaugh JA. Transcription factor Nrg1 mediates capsule formation, stress response, and pathogenesis in *Cryptococcus neoformans*. Eukaryot Cell. 2006;5: 1147–56. doi:10.1128/EC.00145-06

72. Singh A, MacKenzie A, Girnun G, Poeta M Del. Analysis of sphingolipids, sterols, and phospholipids in human pathogenic *Cryptococcus* strains. J Lipid Res. 2017;58: 2017–2036. doi:10.1194/jlr.M078600

73. Mandala SM, Thornton RA, Frommer BR, Curotto JE, Rozdilsky W, Kurtz MB, et al. The discovery of australifungin, a novel inhibitor of sphinganine N-acyltransferase from *Sporormiella australis*. Producing organism, fermentation, isolation, and biological activity. J Antibiot (Tokyo). 1995;48: 349–56. doi:10.7164/antibiotics.48.349

74. Bligh EG, Dyer WJ. A rapid method of total lipid extraction and purification. Can J Biochem Physiol. 1959;37: 911–917. doi:10.1139/o59-099

